# The *Piks* allele of the NLR immune receptor *Pik* breaks the recognition of *AvrPik* effectors of the rice blast fungus

**DOI:** 10.1101/2022.07.19.500709

**Authors:** Gui Xiao, Wenjuan Wang, Muxing Liu, Ya Li, Jianbin Liu, Marina Franceschetti, Zhaofeng Yi, Xiaoyuan Zhu, Zhengguang Zhang, Guodong Lu, Mark J. Banfield, Jun Wu, Bo Zhou

**Affiliations:** State Key Laboratory of Hybrid Rice, Hunan Hybrid Rice Research Center, Changsha 410128, China; International Rice Research Institute, DAPO Box 7777, Metro Manila 1301, Philippines; Guangdong Provincial Key Laboratory of High Technology for Plant Protection, Plant Protection Research Institute, Guangdong Academy of Agricultural Sciences, Guangzhou, 510640, China; Department of Plant Pathology, College of Plant Protection, Nanjing Agricultural University, and Key Laboratory of Integrated Management of Crop Diseases and Pests, Ministry of Education, Nanjing 210095, China; State Key Laboratory of Ecological Pest Control for Fujian and Taiwan Crops, Fujian Agriculture and Forestry University, Fuzhou 350002, China; Department of Biochemistry and Metabolism, John Innes Centre, Norwich Research Park, Norwich, NR4 7UH, UK

## Abstract

Arms race co-evolution of plant-pathogen interactions evolved sophisticated recognition mechanisms between host immune receptors and pathogen effectors. Different allelic haplotypes of an immune receptor in host mount distinct recognition against sequence or non-sequence related effectors in pathogens. We report the molecular characterization of the *Piks* allele of the rice immune receptor *Pik* against rice blast pathogen, which requires two head-to-head arrayed nucleotide binding site and leucine-rich repeat proteins. Like other *Pik* genes, both *Piks*-1 and *Piks*-2 are necessary and sufficient for *Piks*-mediated resistance. However, unlike other *Pik* alleles, *Piks* does not recognize any known *AvrPik* variants of *M. oryzae*. Sequence analysis of the genome of an avirulent isolate V86010 further revealed that its cognate avirulence (*Avr*) gene most likely has no significant sequence similarity to known *AvrPik* variants. We conclude that *Piks* breaks the canonical *Pik*/*AvrPik* recognition pattern. *Piks*-1 and *Pikm*-1 have only two amino acid differences within the integrated heavy metal-associated (HMA) domain. Pikm-1-HMA interact with AvrPik-A, -D and -E in *vitro* and in *vivo*, whereas Piks-1-HMA does not bind any AvrPik variants. Reciprocal exchanges of single amino acid residues between Piks-1 and Pikm-1 further reveal a dynamic recognition mechanism between *Piks/Pikm* alleles and their respective effectors. *Piks-1*^*E229Q*^*/Pikm-1*^*V261A*^ can only activate immunity to *AvrPik-D* but not to other effectors, indicating that the amino acid change of E to Q at position 229 leads to its gain of a partial recognition spectrum of *Pikm*. By contrast, *Piks*-1^A261V^/*Pikm*-1^Q229E^ confers immunity to the *Piks* cognate effector, indicating that the amino acid change of Q to E at position 229 leads to its shifts of the recognition from *Pikm* to *Piks*. Intriguingly, binding activities in both Y2H and analytical gel filtration assays are illustrated between Piks-1^A261V^/Pikm-1^Q229E^ and AvrPik-D. However, it is unable to mount immunity against *AvrPik-D*, suggesting that biochemical activities based on in *vitro* and in *vivo* assays could be insufficient for sustaining biological function of receptor and effector pairs.

## INTRODUCTION

Pathogen/microbe-associated molecular patterns (PAMPs/MAMPs) are recognized by host cell surface-localized pattern-recognition receptors (PRRs) to active the first layer of plant immunity (Jones and Dangl 2006; Zhang and Zhou 2010). Pathogens deliver a myriad of effectors into host cells to inhibit such host immune responses and /or create a favorable host cell environment for infection (Li et al. 2017). Plants have developed cytoplasmic immune receptors of the Nucleotide-binding domain leucine-rich repeat class (NBS-LRRs, known as NLRs), which directly or indirectly recognize pathogen effectors leading to intracellular immunity (Jones and Dangl 2006). This layer of immunity confers strong resistance, often associated with localised cell death known as the hypersensitive response (HR). Most plant NLRs are defined by a multi-domain architecture with central nucleotide-binding (NB-ARC) and C-terminal LRR regions, and either an N-terminal TOLL/interleukin-1 receptor (TIR) or a coiled-coil (CC) domain (Jones and Dangl 2006; Takken and Goverse 2012).

Rice blast disease, caused by the fungal pathogen *Magnaporthe oryzae* (*M. oryzae*), is the most devastating disease threating global rice production and food security (Skamnioti and Gurr 2009; Asibi et al. 2019; Shahriar et al. 2020). The most effective and environmental-friendly approach to controlling rice blast is the deployment of rice blast resistance (*R*) gene. To date, more than 30 rice blast *R* genes have been characterized, most of which encode NLRs, with the exception of *Pid2*, *Pi21, and Pita2/Ptr. Pid2* encodes a β-lectin receptor-like kinase (Chen et al. 2006); *Pi21* encodes a proline-rich protein (Fukuoka et al. 2009); and *Pita2*/*Ptr* encodes an atypical *R* protein with four Armadillo repeats (Zhao et al. 2018; Meng et al. 2020). However, only a limited number of *M. oryzae* avirulence (*Avr*) genes (also known as effectors), recognized by these *R* genes have been cloned to date. They include *AvrPita* (Orbach et al. 2000), *AvrPia* (Yoshida et al. 2009), *AvrPik* and its alleles (Yoshida et al. 2009), *AvrPii* (Yoshida et al. 2009), *AvrPiz-t* (Li et al. 2009), *Avr1-CO39* (Ribot et al. 2013), *AvrPib* (Zhang et al. 2015), and *AvrPi9* (Wu et al. 2015). Based on the pattern by which rice blast *R* genes recognize their respective *Avr* genes, distinct mechanisms can be described for their function. Pairs of *Pita/AvrPita* and *Pib/AvrPib* represent the first mode in which a single *R* gene recognizes a single *Avr* gene (Jia et al. 2000; Zhang et al. 2015). Pairs of *Pi-CO39*/*Avr1-CO39* and *Pia*/*AvrPia* represent the second mode in which a single *R* gene recognizes two sequence-unrelated *Avr* genes (Yoshida et al. 2009; Okuyama et al. 2011; Cesari et al. 2013). Pairs of *Pi9*/*AvrPi9* and *Piz-t*/*AvrPiz-t* represent the third mode in which different *R* alleles each recognize sequence-unrelated *Avr* genes (Wu et al. 2015; Li et al. 2009). Pairs of *Pik* alleles and *AvrPik* variants represent the fourth mode in which different *R* alleles have distinct recognition specificities to different *Avr* variants (Yoshida et al. 2009; Kanzaki et al. 2012).

Molecular mechanisms underlying recognition of *AvrPik* variants by *Pik* alleles have been extensively characterized. Both the *Pik-1*/*Pik-2* NLR pair and *AvrPik* effectors exist in an allelic series in rice and *M. oryzae* populations. The *Pik* locus is located on rice chromosome 11, where at least nine genes, *Pik*, Pikm, Pikp, Pikh, Pi1, Pi7, Pik, Piks* and *Pike*, have been identified (Ashikawa et al. 2012; Ashikawa et al. 2008; Yuan et al. 2011; Zhai et al. 2014; Hua et al. 2012; Campbell et al. 2004; Zhai et al. 2011; Chen et al. 2015). For *AvrPik*, six different variants (A–F) have been identified with 1–3 amino acid differences between pathogen isolates (Yoshida et al. 2009; Kanzaki et al. 2012; Wu et al. 2014; Longya et al. 2019). The direct interaction between *Pik* and *AvrPik* is mediated by the HMA domain of *Pik-1*, with this domain being integrated between the coiled-coil and nucleotide-binding (NB-ARC) domains (Maqbool et al. 2015; De la Concepcion et al. 2018, 2021). At the sequence level, *Pikp-1* and *Pikm-1* predominantly differ in their HMA domains, and this underpins different effector recognition specificities. *Pikp* only recognizes *AvrPik*-D, whereas *Pikm* can recognize *AvrPik-*D, *AvrPik*-A and *AvrPik*-E (Kanzaki et al. 2012; De la Concepcion et al. 2018). The interface between *AvrPik* effectors and the HMA domain of both *Pikp* and *Pikm* has been structurally characterized (Maqbool et al. 2015; De la Concepcion et al. 2018). The *AvrPik*-C and *AvrPik*-F effector variants are not currently recognized by any *Pik* alleles (Kanzaki et al. 2012; Longya et al. 2019). However, the *AvrPik-C* effector variant has been reported to interact with *Pikh*-HMA *in vitro*, with sufficient affinity to allow crystallization of the complex (De la Concepcion et al. 2021). Recently, the potential to engineer new-to-nature receptors specificities, such as recognition of stealthy effector variants has been demonstrated, which has broad implications for rational design of plant NLRs (De la Concepcion et al. 2019; De la Concepcion et al. 2021; Maidment et al. 2022).

The *Piks* gene was identified in the monogenic line IRBLKs-F5, and originated from the donor variety Fujisaka 5. It is responsible for the resistance against the rice blast isolate V86010 (Tsunematsu et al. 2000). However, the mechanism underlying this resistance has remained elusive, and the specificity of interaction with known *AvrPik* variants has not been deciphered. In this study, we conduct molecular characterization of *Piks* through genetic analyses and interaction assays with different *AvrPik* variants. The function of key amino acids differing between Piks and Pikm is investigated to provide insights into the evolution of distinct specificities among different *Pik* alleles.

## RESULTS

### *Piks* in IRBLKs-F5 confers novel race-specific resistance against rice blast

To verify that *Piks* identified in the monogenic line IRBLks-F5 confers race-specific resistance against rice blast, the response of IRBLks-F5 towards pathogen isolates V86010 and CA89 was evaluated. As shown in Figure 1A, IRBLKs-F5 is resistant to V86010, but susceptible to other isolates. Its recipient rice variety Lijiangxintuanheigu (LTH) is susceptible to all isolates. These results illustrate that *Piks* in IRBLKs-F5 determines a race specific resistance to rice blast. A set of 367 rice blast isolates from the Philippines was further used to evaluate and compare the resistance spectra conferred by different *Pik* alleles (Table S1). As Table S1 shows, the *Piks* monogenic line, IRBLks-F5 was susceptible to all isolates except V86010, showing a 0.27% of resistance frequency to the tested isolates (Figure 1B). By contrast, other *Pik* monogenic lines showed varying resistance frequencies ranging from 42.2% for IRBLK-Ka and IRBL7-M to 73.8% for IRBLKp-K60 (Figure 1B). It is worth noting that the higher resistance frequency of IRBLKp-K60 is unlikely to be only determined by *Pikp* due to the presence of an additional *R* gene linked to the *Pi19* locus, as previously characterized (Selisana et al. 2017). These results demonstrate that *Piks* controls a much narrower resistance spectrum than other *Pik* alleles against rice blast isolates in the Philippines.

**Figure 1.**
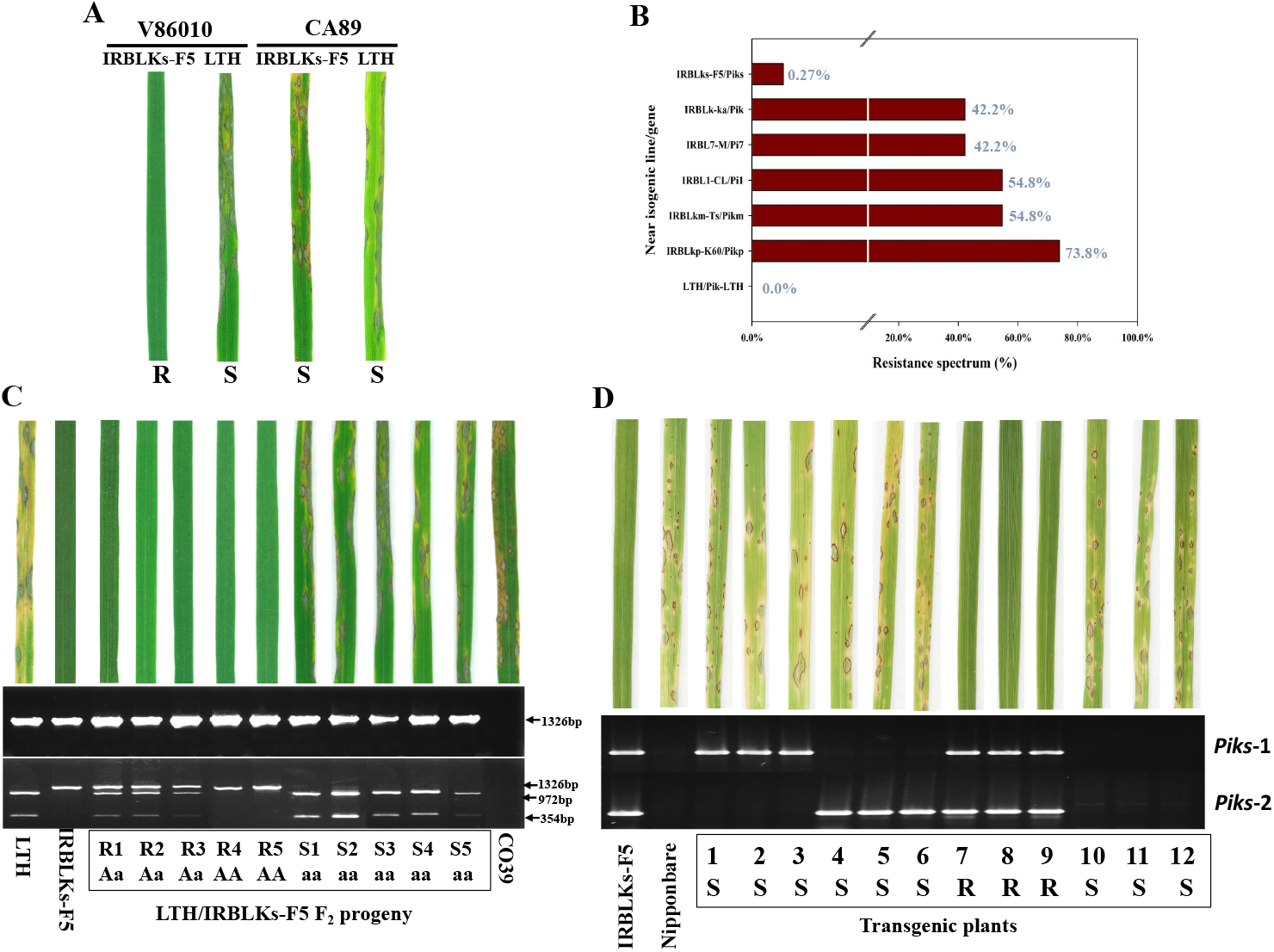
Characterization and cloning of *Piks*. **(A)** Identification of a *M. oryzae* isolate avirulence to *Piks* gene near isogenic line IRBLKs-F5. V86010 and CA89 are two *M. oryzae* isolates used for inoculation. **(B)** Resistance spectrum of *Piks* and other *Pik* allelic genes. LTH (Lijiangxintuanheigu) is the susceptible control for inoculation and also the recipient cultivar for the development of monogenic line. **(C)** Co-segregation analysis of *Piks*-mediated resistance. Reaction to inoculation with the blast isolate V86010 in CO39, LTH, IRBLKs-F5 and a set of F_2_ progeny. PCR amplification assay of the parental lines and F_2_ progeny using primer pair RGA4-F3/RGA4-R3. Enzymatic digestion results of the PCR products using *PstI*. R1 to R5 means the resistance progeny 1 to 5, S1 to S5 means the susceptible progeny 1 to 5. AA homozygous resistant, Aa heterozygote, aa homozygous susceptible. **(D)** Complementation testing and molecular analysis of the transgenic plants. Reaction of IRBLKs-F5, Nipponbare, and a set of F_2_ progeny (numbers 1–12) derived from a cross between T1 plants harboring *Piks*-1 and *Piks*-2, individually, to the blast isolate V86010. The isolate V86010 is avirulence to the *Piks* donor cultivar IRBLKs-F5 and virulence to the transformation recipient cultivar Nipponbare. Co-segregation of the resistance phenotype with the presence of both the *Piks*-1 gene and the *Piks*-2 gene. The transgenic lines numbered 1 to 3 harbor only *Piks*-1, those numbered 4 to 6 contain only *Piks*-2, number 7-9 contain both *Piks*-1 and *Piks*-2, and that numbered 10-12 harbor neither of the transgenes.

To further investigate whether *Piks* resistance is dependent on the recognition of *AvrPik*, a set of transformed pathogen isolates, using the recipient isolate R88-002 that does not harbor *AvrPik*, was generated with each containing one of the six known *AvrPik* variants (*AvrPik*-*A* to -*F*) (Figure S1). Pathogenicity tests against different *Pik* monogenic lines were then performed. As shown in Table 1, IRBLKs-F5 was susceptible to all transformed isolates. However, *Pikp, Pi7*, and *Pik* monogenic lines were resistant to *AvrPik*-*D* transformed isolates; *Pikm* and *Pi1* monogenic lines were avirulent to *AvrPik-A*, -*D*, and -*E* transformed isolates (Table 1). This *R*-*Avr* interaction assay demonstrates that *Piks* was unable to activate rice immunity against any of the six known *AvrPik* variants.

**Table 1.** Disease reactions of different *Pik* alleles in the near isogenic lines to *M. oryzae* R88-002 isolate expressing six *AvrPik* variants.

| Strains | *AvrPik* variants | Differential near isogenic lines and respective *Pik* alleles |  |  |  |  |  | LTH |
| --- | --- | --- | --- | --- | --- | --- | --- | --- |
|  |  | IRBLKs-F5 | IRBL7-M | IRBLKp-K60 | IRBLK-Ka | IRBLKm-Ts | IRBL1-CL |  |
|  |  | *Piks* | *Pi7* | *Pikp* | *Pik* | *Pikm* | *Pil* |  |
| R88-002<sup>*AvrPik-A*</sup>-1 | *AvrPik-A* | S | S | S | S | R | R | S |
| R88-002<sup>*AvrPik-A*</sup>-25 | *AvrPik-A* | S | S | S | S | R | R | S |
| R88-002<sup>*AvrPik-B*</sup>-3 | *AvrPik-B* | S | S | S | S | S | S | S |
| R88-002<sup>*AvrPik-B*</sup>-4 | *AvrPik-B* | S | S | S | S | S | S | S |
| R88-002<sup>*AvrPik-C*</sup>-3 | *AvrPik-C* | S | S | S | S | S | S | S |
| R88-002<sup>*AvrPik-C*</sup>-14 | *AvrPik-C* | S | S | S | S | S | S | S |
| R88-002<sup>*AvrPik-D*</sup>-4 | *AvrPik-D* | S | R | R | R | R | R | S |
| R88-002<sup>*AvrPik-D*</sup>-17 | *AvrPik-D* | S | R | R | R | R | R | S |
| R88-002<sup>*AvrPik-E*</sup>-1 | *AvrPik-E* | S | S | S | S | R | R | S |
| R88-002<sup>*AvrPik-E*</sup>-10 | *AvrPik-E* | S | S | S | S | R | R | S |
| R88-002<sup>*AvrPik-F*</sup>-1 | *AvrPik-F* | S | S | S | S | S | S | S |
| R88-002<sup>*AvrPik-F*</sup>-14 | *AvrPik-F* | S | S | S | S | S | S | S |
| R88-002 | None | S | S | S | S | S | S | S |

R88-002 is the recipient rice blast isolate; R, resistance; S, susceptible

### Both *Piks*-1 and *Piks*-2 are required for *Piks*-mediated resistance

To determine whether *Piks* is responsible for the resistance of IRBLKs-F5 against V86010, an F_2_ population was developed from a cross between IRBLKs-F5 and its recipient rice variety Lijiangxintuanheigu (LTH). A total of 1387 individuals of the F_2_ population were inoculated with V86010. As indicated in Table 2, the F_2_ population segregated for 1053 resistant and 334 susceptible plants, showing an expected 3:1 resistance versus susceptibility ratio [*χ*^2^(3:1) = 0.625]. This result confirms that the resistance of IRBLks-F5 against V86010 is controlled by a single gene or genetic locus.

**Table 2.**
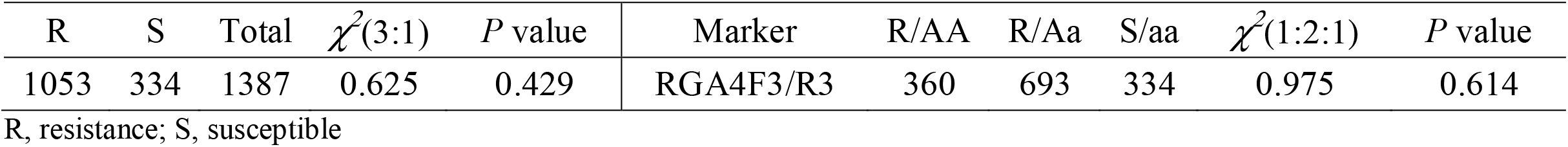
Genetic and co-segregation analysis of the LTH/IRBLKs-F5 F_2_ population inoculated with the rice blast isolate V86010

| R | S | Total | $\chi^2(3:1)$ | *P* value | Marker | R/AA | R/Aa | S/aa | $\chi^2(1:2:1)$ | *P* value |
| --- | --- | --- | --- | --- | --- | --- | --- | --- | --- | --- |
| 1053 | 334 | 1387 | 0.625 | 0.429 | RGA4F3/R3 | 360 | 693 | 334 | 0.975 | 0.614 |

R, resistance; S, susceptible

We cloned and determined the sequences of both *Piks-1* and *Piks-2* in IRBLKs-F5, and these were identical to those deposited in NCBI (Genbank accession no.: HQ662329). A gene specific molecular marker RGA4-F3/R3 showing polymorphism between IRBLKs-F5 and LTH was used for the molecular analysis of the F_2_ progeny (Figures 1C, S2). We found that all resistant progeny exhibited marker pattern of the resistant parent (and all susceptible progeny exhibited the susceptible parent pattern), indicating that *Piks* co-segregates with RGA4-F3/R3 (Figure 1C; Table 2). To determine whether the function of *Piks* requires co-expression of both *Piks*-1 and *Piks*-2, *Piks*-1/*Piks*-2 transgenic rice plants were generated via crossing and selfing of *Piks*-1 and *Piks*-2 individual transgenic plants. Analysis of transgenic plants showed that only *Piks*-1 and *Piks*-2 combined transgenic lines showed resistance against V86010 (Figure 1D). By contrast, their donor transgenic lines with only either *Piks*-1 or *Piks*-2 were susceptible (Figure 1D). These results confirmed genetically that the *Piks* resistance requires co-expression of both *Piks*-1 and *Piks*-2. In addition, we generated three independent knockout mutants of *Piks*-1 and *Piks*-2 using CRISPR. As illustrated in Figure 2A, each of the knockout mutants Piks-KO-3, -4, and -7 had a distinct mutation pattern in either *Piks*-1 or *Piks*-2. It is noted that Piks-KO-3 and Piks-KO-7 had deletions of 6- and 30-nucleotide in *Piks*-1, respectively, whereas they had the same single-nucleotide insertion in *Piks*-2 (Figure 2A). Based on the frameshift mutations in *Piks*-1, *Piks*-2, or both, we speculated that the function of either *Piks*-1 or *Piks*-2 was likely disrupted in three mutants. The resistance phenotype of these mutants to V86010 was investigated, revealing that they were compromised (Figure 2B), further verifying that both *Piks*-1 and *Piks*-2 are required for the *Piks*-mediated resistance.

**Figure 2.**
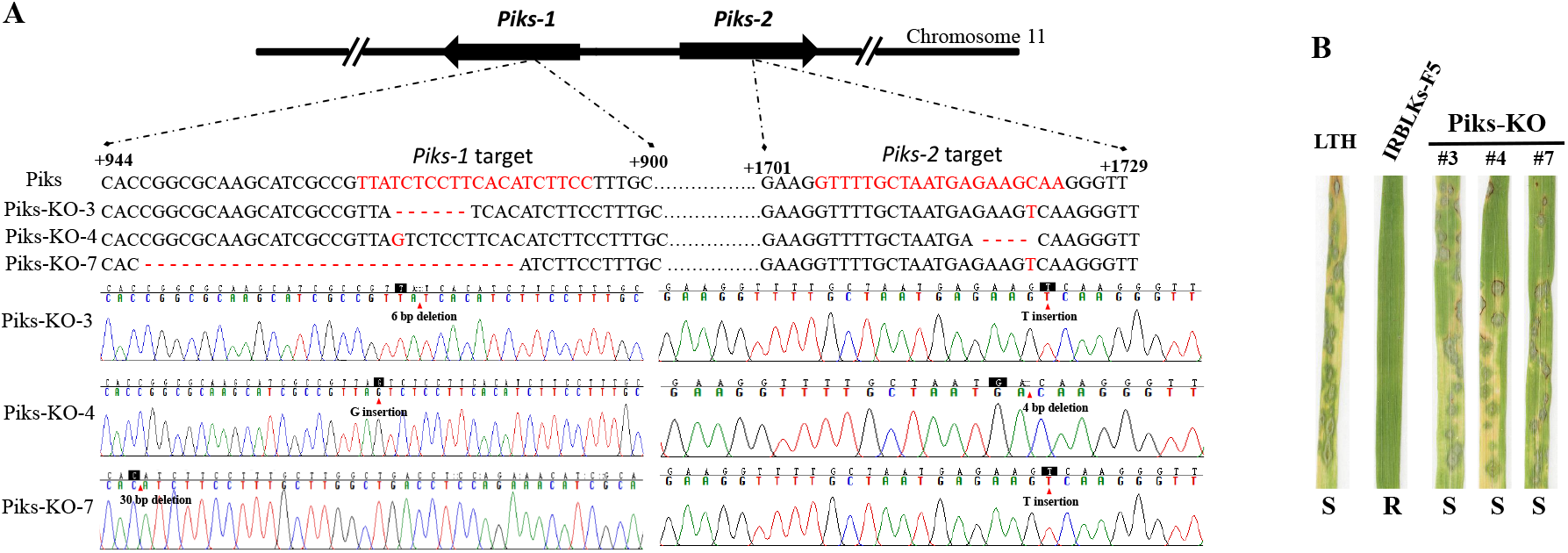
Genotype and phenotypes of *Piks* loss-of-function mutants created by CRISPR technology. **(A)** Schematic description of the CRISPR mutant of *Piks*-1 and *Piks*-2. Mutations detected in genomic DNA of *Piks*-KO-3, *Piks*-KO-4 and *Piks*-KO-7 mutants are shown by SANGER sequencing chromatography. (B) Phenotype of *Piks* loss-of-function mutants. Rice blast isolate V86010 is used for inoculation. LTH (Lijiangxintuanheigu) and IRBLKs-F5 are used as control.

### *Piks*-mediated resistance against V86010 is not determined by *AvrPik*

The V86010 strain contains two *AvrPik* variants, *AvrPik*-*D* and *-E*, as determined via sequencing of PCR amplicons (Figure S3). The presence of *AvrPik* variants was further investigated through homology search against the V86010 whole-genome-shotgun (WGS) sequence (ASM210529v1). Only a single WGS contig (MWIT01000852.1) having over 99.7% sequence identity was identified, which was further found to derive from an assembly of both *AvrPik*-*D* and *-E* sequences due to extreme sequence similarity. Therefore, we conclude V86010 only contains two *AvrPik* variants. To investigate whether *AvrPik-D* and *-E* is involved in recognition of V86010 by *Piks*, V86010 mutant strains disrupted in both *AvrPik-D* and *-E* were generated and used in pathogenicity assays. Out of 268 hygromycin resistant transformed strains, three (#3, #4, and #38) were found to be true knockouts of both *AvrPik*-*D* and -*E* as they displayed none of the expected amplicons of *AvrPik* whereas they all contained the DNA fragment of the selectable marker (hygromycin B phosphotransferase gene) flanked by the homology arm (Figure S4). Pathogenicity tests revealed that these three mutant strains were still recognized by IRBLks-F5 which retains a resistant phenotype (Figure 3A). In addition, they resulted in gain-of-susceptiblity to other *Pik* monogenic lines (Figure 3A), verifying previous findings of *AvrPik* and *Pik* interactions (Kanzaki et al. 2012). Finally, these three mutant strains were used to challenge the transgenic *Piks* lines described above. All transgenic *Piks* lines harboring both *Piks-1* and *Piks-2* were resistant to these three mutant strains (Figure 3B). Taken together, these results suggest that V86101 contains a currently unknown *Avr* gene recognized by *Piks*, leading to activation of immunity. Moreover, this novel *Avr* gene cognate to *Piks* most likely has no significant sequence similarity to known *AvrPik* alleles.

**Figure 3.**
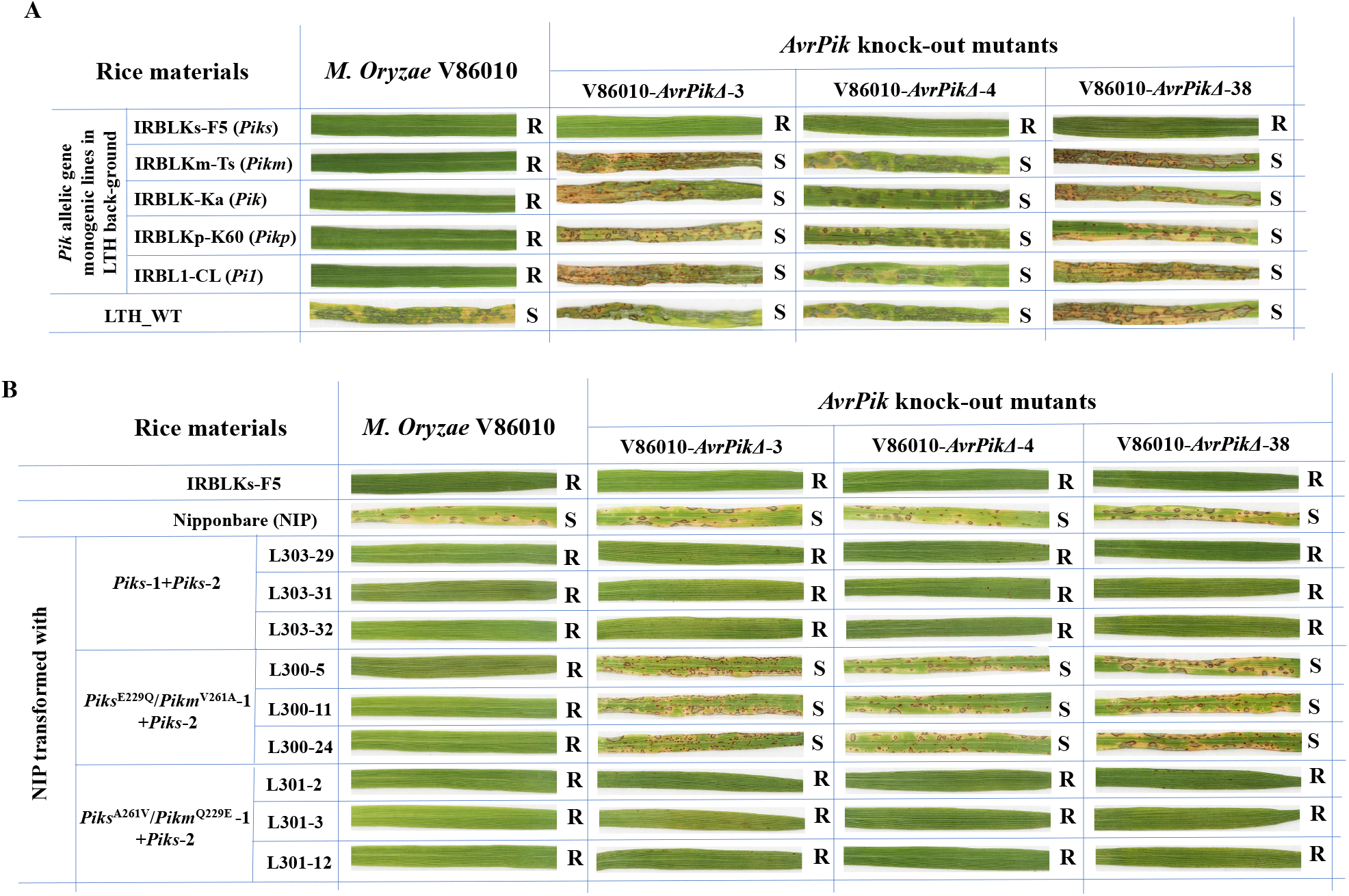
The avirulence of V86010 to *Piks* do not rely on *AvrPik*. (A) Disease reaction of the *Pik* allelic gene monogenic lines inoculated with V86010 and three AvrPik-D/-E knock-out mutant strains. (B) Disease reaction of the *Piks* and *Piks* single amino acid substitution mutant (*Piks*^E229Q^/*Pikm*^V261A^ and *Piks*^A261V^/*Pikm*^Q229E^) transgenic plants challenged with blast isolate V86010 and V86010 knout-out of *AvrPik*-D/-E strains. NIP is the recipient variety of the transgenes.

### Two amino acids that distinguish Piks-1 from Pikm-1 are critical for resistance specificities

It is well established that *Pik*-1 alleles have a higher level of sequence variation than the *Pik*-2 alleles (Figures S5, S6). It is worth noting that the *Piks*-2 sequence is identical to *Pikm*-2 and *Pik*-2, suggesting that they are unlikely be the determinants of differential recognition specificities (Figure S6). Nevertheless, *Piks*-1 is most sequence related to *Pikm*-1 and *Pi1*-5 (Figure 4A, Figure S5). There are two amino acids differing *Piks*-1 from *Pikm*-1 and *Pi1*-5 at amino acid positions of 229 and 261. In *Piks*-1, position 229 is a glutamate residue (E) whilst in *Pi1*-5 it is an aspartate (D) and in *Pikm*-1 a glutamine (Q). At position 261, the residues are Alanine (A) for *Piks*-1 and Valine (V) for both *Pi1*-5 and *Pikm*-1 (Figure 4A, Figure S5). Given the amino acid sequences of Piks-2 and Pikm-2 are identical, we focused on the characterization of amino acid differences between these two haplotypes.

**Figure 4.**
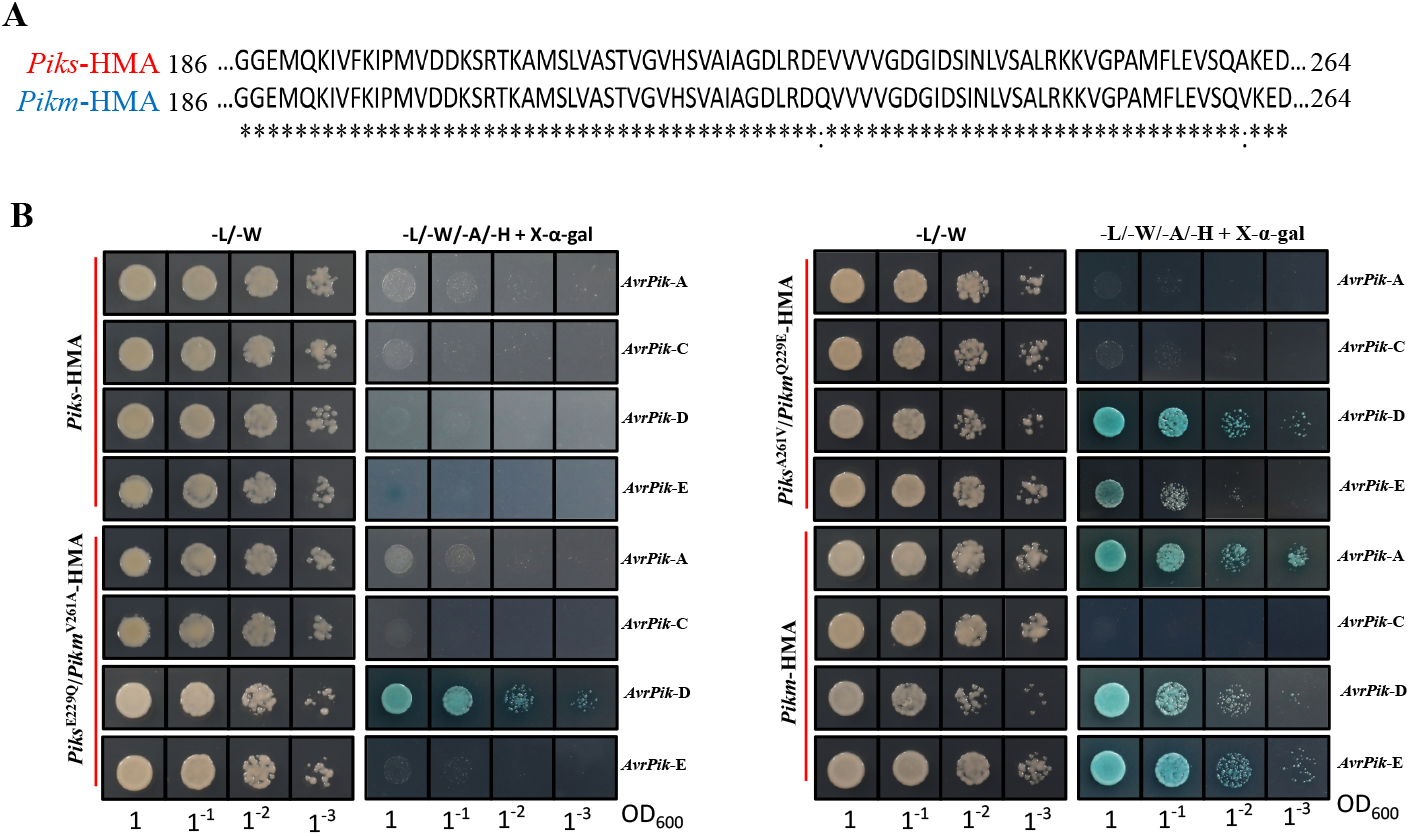
Yeast two hybrid assay demonstrate the binding of *AvrPik* effector variants to the HMA domain of *Piks, Pikm* and *Piks* single amino acid substitution mutants (*Piks*^E229Q^/*Pikm*^V261A^ and *Piks*^A261V^/*Pikm*^Q229E^). (A) Amino acid sequence alignment of *Piks*-1 and *Pikm*-1 HMA domains. (B) *Piks-*HMA does not interaction with any *AvrPik* variants as assayed by Y2H. The control plate for yeast growth is on the left, with the selective plate on the right. Each experiment was repeated a minimum of three times, with similar results.

To enable genetic characterization in rice, two variants of *Piks-1*, introducing the amino acid residue at the equivalent position in Pikm-1, which are *Piks*-1^E229Q^ and *Piks*-1^A261V^, were generated and used for transformation of rice cultivar Nipponbare (Nipponbare was also transformed with *Piks*-1*/Piks*-2 as a control). The resulting transgenic plants were challenged with the *M. oryzae* strain V86010 and V86010 knockouts in *AvrPik*-*D/E*. The *Piks*-1^E229Q^*/Piks*-2 transgenic plants were resistant to V86010, the same phenotype observed for the *Piks*-1*/Piks*-2 transgenic plants (Figure 3B). By contrast, we found that *Piks*-1^E229Q^*/Piks*-2 transgenic plants were susceptible to all the *AvrPik*-D/E-knockout mutant strains (Figure 3B). Therefore, we deduced from this data that the switch from E to Q at position 229 leads to a loss of *Piks*-mediated resistance against the cognate (as yet unknown) *Avr* gene. To investigate which *AvrPik* haploptype is responsible to *Piks*-1^E229Q^/*Piks*-2, the resistance phenotypes were assessed using the same set of R88-002 transformed isolates containing different *AvrPik* haplotypes described above. As shown in Table 3, *Piks*-1^E229Q^/*Piks*-2 transgenic plants were only resistant to *AvrPik*-*D* containing R88-002 strains, and not other avirulent haplotypes, demonstrating a distinct spectrum compared to *Pikm* that recognizes *AvrPik*-*D*, -*E*, and -*A*. The *Piks*-1^E229Q^ mutation generates a protein equivalent to a *Pikm*-1^V261A^ mutation. Therefore, our data can also be interpreted as the switch from V to A at the position 261 leads to a loss of *Pikm* recognition to both *AvrPik*-*E* and -*A*, but retains the recognition of *AvrPik*-*D* in transgenic rice (Table 3).

**Table 3.**
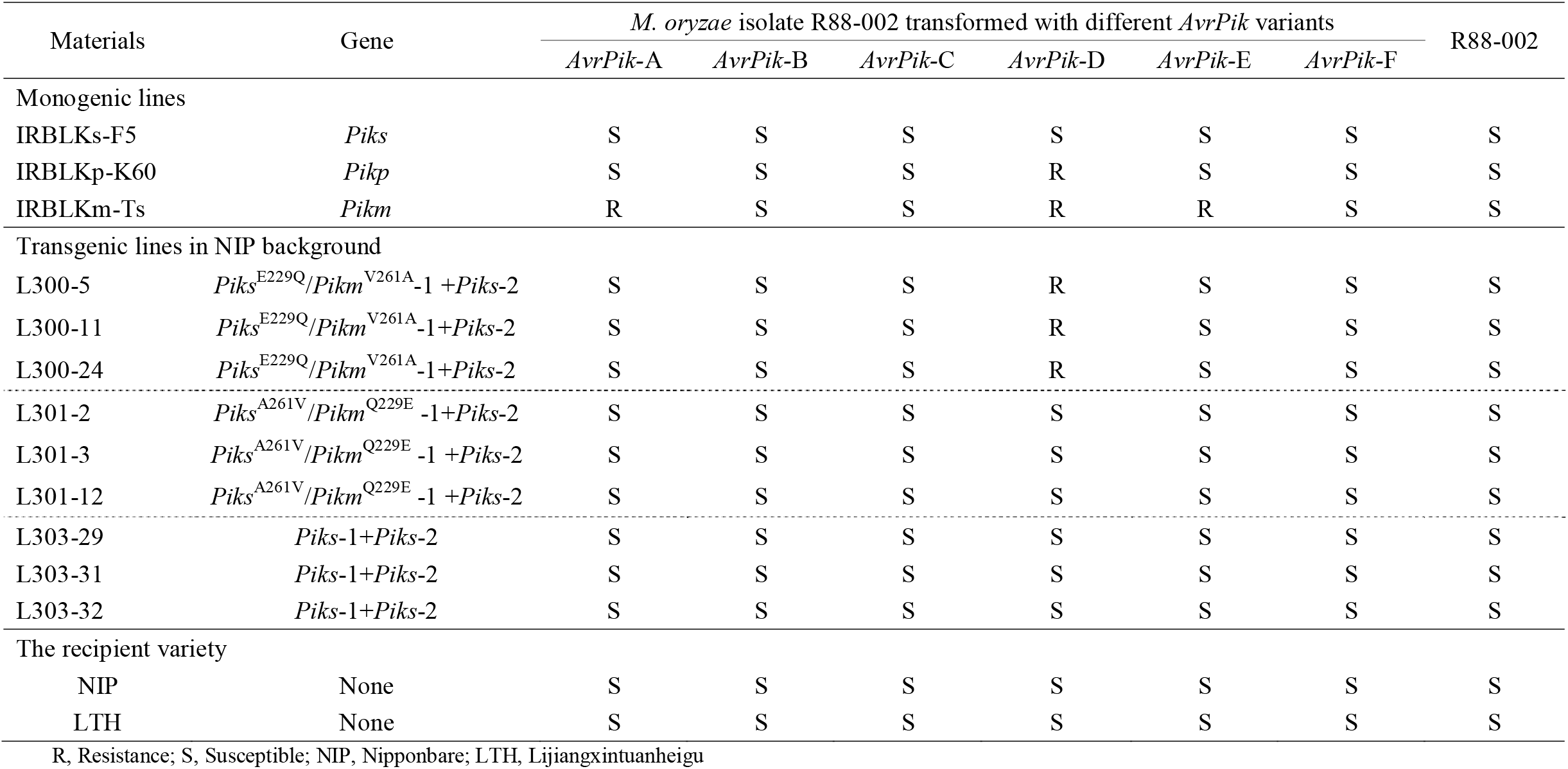
Disease reactions of *Piks* gene and two *Piks* mutants *Piks*^E229Q^*/Pikm*^V261A^ and *Piks*^A261V^*/Pikm*^Q229E^ transgenic lines to *M. oryzae* R88-002 isolate expressing six *AvrPik* variants.

| Materials | Gene | *M. oryzae* isolate R88-002 transformed with different *AvrPik* variants |  |  |  |  |  | R88-002 |
| --- | --- | --- | --- | --- | --- | --- | --- | --- |
|  |  | *AvrPik*-A | *AvrPik*-B | *AvrPik*-C | *AvrPik*-D | *AvrPik*-E | *AvrPik*-F |  |
| Monogenic lines |  |  |  |  |  |  |  |  |
| IRBLKs-F5 | *Piks* | S | S | S | S | S | S | S |
| IRBLKp-K60 | *Pikp* | S | S | S | R | S | S | S |
| IRBLKm-Ts | *Pikm* | R | S | S | R | R | S | S |
| Transgenic lines in NIP background |  |  |  |  |  |  |  |  |
| L300-5 | *Piks*<sup>E229Q</sup>/*Pikm*<sup>V261A</sup>-1 +*Piks*-2 | S | S | S | R | S | S | S |
| L300-11 | *Piks*<sup>E229Q</sup>/*Pikm*<sup>V261A</sup>-1+*Piks*-2 | S | S | S | R | S | S | S |
| L300-24 | *Piks*<sup>E229Q</sup>/*Pikm*<sup>V261A</sup>-1+*Piks*-2 | S | S | S | R | S | S | S |
| L301-2 | *Piks*<sup>A261V</sup>/*Pikm*<sup>Q229E</sup>-1+*Piks*-2 | S | S | S | S | S | S | S |
| L301-3 | *Piks*<sup>A261V</sup>/*Pikm*<sup>Q229E</sup>-1 +*Piks*-2 | S | S | S | S | S | S | S |
| L301-12 | *Piks*<sup>A261V</sup>/*Pikm*<sup>Q229E</sup>-1 +*Piks*-2 | S | S | S | S | S | S | S |
| L303-29 | *Piks*-1+*Piks*-2 | S | S | S | S | S | S | S |
| L303-31 | *Piks*-1+*Piks*-2 | S | S | S | S | S | S | S |
| L303-32 | *Piks*-1+*Piks*-2 | S | S | S | S | S | S | S |
| The recipient variety |  |  |  |  |  |  |  |  |
| NIP | None | S | S | S | S | S | S | S |
| LTH | None | S | S | S | S | S | S | S |

R, Resistance; S, Susceptible; NIP, Nipponbare; LTH, Lijiangxintuanheigu

We also challenged the *Piks-*1^A261V^*/Piks*-2 transgenic Nipponbare plants against V86010 and with its knockouts of *AvrPik*-*D/E* and found they were resistant to all strains (Figure 3B). This result suggests that the mutation from A to V at the position 261 does not compromise the function of *Piks* to the cognate (as yet unknown) *Avr* gene. Further, they were susceptible to R88-002 transformed strains with the six *AvrPik* variants (Table 3). Likewise, *Piks*-1^A261V^ can be designated as *Pikm*-1^Q229E^. Therefore, we deduced that the switch from Q to E at position 229 results in the loss of function of *Pikm* to *AvrPik*-*D*, -*E*, and -*A*.

### Piks-1, Pikm-1, Piks-1^E229Q^/Pikm-1^V261A^, and Piks-1^A261V^/Pikm-1^Q229E^ differentially interact with known AvrPik proteins

To investigate the interaction pattern of *Piks* and its alleles with known *AvrPik* variants, a protein-protein binding assay using a yeast two-hybrid (Y2H) was conducted. As shown in Figure 4B, no significant yeast growth on selective media was observed when the HMA domain of Piks (Piks-HMA) was co-transformed with each of four AvrPik effector variants (AvrPik-A, -C, -D and -E). By contrast, significant yeast growth was observed when the HMA domain of Pikm (Pikm-HMA) co-transformed with *AvrPik*-*A, AvrPik*-*D*, and *AvrPik*-*E*, but not *AvrPik*-*C* (Figure 4B). β-galactosidase activity assay, resulting in the development of blue coloration, further verified the binding activity between each interacting pair. These results from Pikm-HMA are consistent with previous findings reported by different laboratories (Kanzaki et al. 2012; De la Concepcion et al. 2018). Therefore, we speculate that these two amino acid residues differentiating Pikm from Piks are critical for binding activity between Pikm and its cognate AvrPik variants.

Next, we tested the interaction of the HMA domains of Piks^E229Q^/Pikm^V261A^ (Piks^E229Q^/Pikm^V261A^-HMA) and Piks^A261V^/Pikm^Q229E^ (Piks^A261V^/Pikm^Q229E^-HMA) with each of four AvrPik effector variants (AvrPik-A, -C, -D and -E). Firstly, for the Piks^E229Q^/Pikm^V261A^-HMA, we only observed yeast growth and development of blue coloration of co-transformants with *AvrPik*-D and not other effector variants (Figure 4B). Secondly, for the Piks^A261V^/Pikm^Q229E^ variant, we observed yeast growth and development of blue coloration of co-transformants with *AvrPik-D* and *-E*. However, the latter is significantly lower in growth than the former (Figure 4B). Unlike *Pikm*, binding activity between Piks^A261V^/Pikm^Q229E^ -HMA and AvrPik-A was not deduced based on the yeast growth (Figure 4B).

We also took advantage of the well-established methods for production of the AvrPik effectors and Pik-HMA domains in vitro to express and purify AvrPik-A, -C, -D, or -E, Pikm-HMA, Piks-HMA, Piks^E229Q^/Pikm^V261A^-HMA and Piks^A261V^/Pikm^Q229E^-HMA proteins and then test for their interactions using an analytical gel filtration assay (Figure 5). Following mixing and incubation of purified proteins, this assay demonstrated complex formation between Pikm-HMA with AvrPik-A, -D, or -E, but not with AvrPik-C, consistent with previous results (De la Concepcion et al. 2018). By contrast, Piks-HMA did not form complexes with any of AvrPik proteins, which is consistent with the Y2H results (Figure 5). For Piks^E229Q^/Pikm^V261A^-HMA and Piks^A261V^/Pikm^Q229E^-HMA, we found that each mutant formed complexes with AvrPik-D but not with other AvrPik variants (Figure 5). Although weak interaction between Piks^A261V^/Pikm^Q229E^-HMA and AvrPik-E was observed in the Y2H assay, it was not detected by analytical gel filtration with purified proteins.

**Figure 5.**
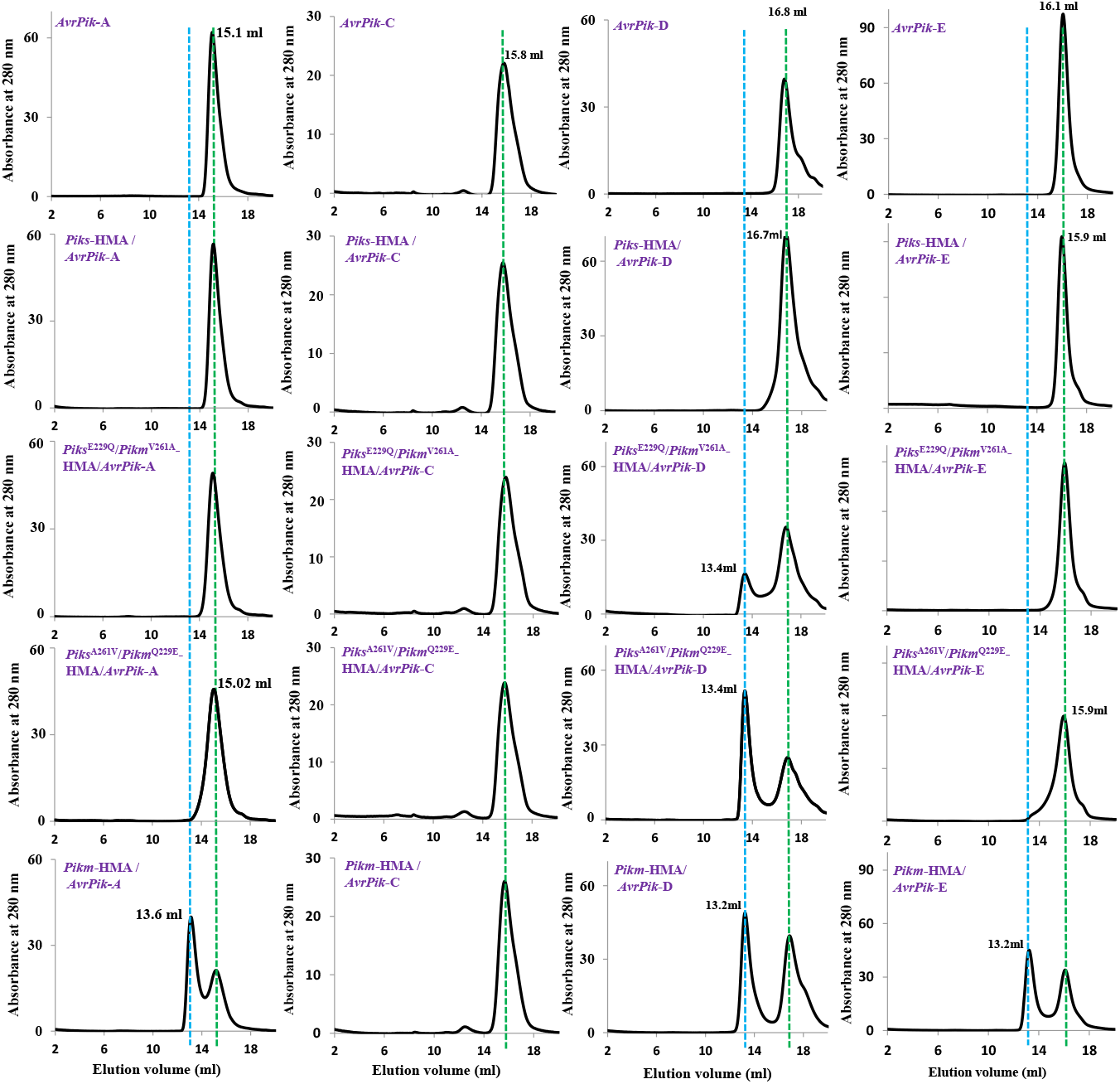
*Piks*-HMA can not binding to any *AvrPik* variants. Analytical Gel Filtration traces depicting the retention volume of *AvrPik* variants (*AvrPik*-A, *AvrPik*-C, *AvrPik*-D and *AvrPik*-E) and *Pik*-HMAs (*Piks*-HMA, *Piks*^E229Q^/*Pikm*^V261A^-HMA and *Piks*^A261V^/*Pikm*^Q229E^-HMA) and the complex. *Pikm*-HMA and the complex are used as the control. Representative SDS-PAGE gels of *AvrPik-*D variants and *AvrPik-D*/*Pikm* protein complex were shown in Figure S7. Each experiment was repeated a minimum of three times, with similar results.

### Piks-1^E229Q^/Pikm-1^V261A^ and Piks-1^A261V^/Pikm-1^Q229E^ elicit weak cell death in N. benthamiana when co-expressed with Piks-2 and AvrPik-D

Finally, we tested the potential of *Piks, Piks-1*^E229Q^/*Pikm-1*^V261^ and *Piks-1*^A261V^/*Pikm-1*^Q229E^ to elicit cell death in *N. benthamiana* leaves (development of necrotic tissue and accumulation of phenolic compounds) with the *AvrPik* variants *AvrPik*-*A*, -*C*, -*D*, or -*E* using *Agrobacterium*-mediated transformation. All constructs were co-expressed with *Piks*-2. Compared to the strong cell death observed in positive control (co-expression of *Pikm* and *AvrPik*-*D*), we observed only moderate cell death for the combinations of *Piks-1*^E229Q^/*Pikm-1*^V261A^+*Piks*-2+*AvrPik*-*D* and *Piks-1*^A261V^/*Pikm-1*^Q229E^+*Piks-2*+*AvrPik*-*D* (Figure 6). For all other combinations, including *Piks* with all effector variants, and *Piks-1*^A261V^/*Pikm-1*^Q229E^ and *Piks-1*^A261V^/*Pikm-1*^Q229E^ with effector variants *AvrPik*-*A, -C*, or -*E* we did not observe signatures of cell death (Figure 6).

**Figure 6.**
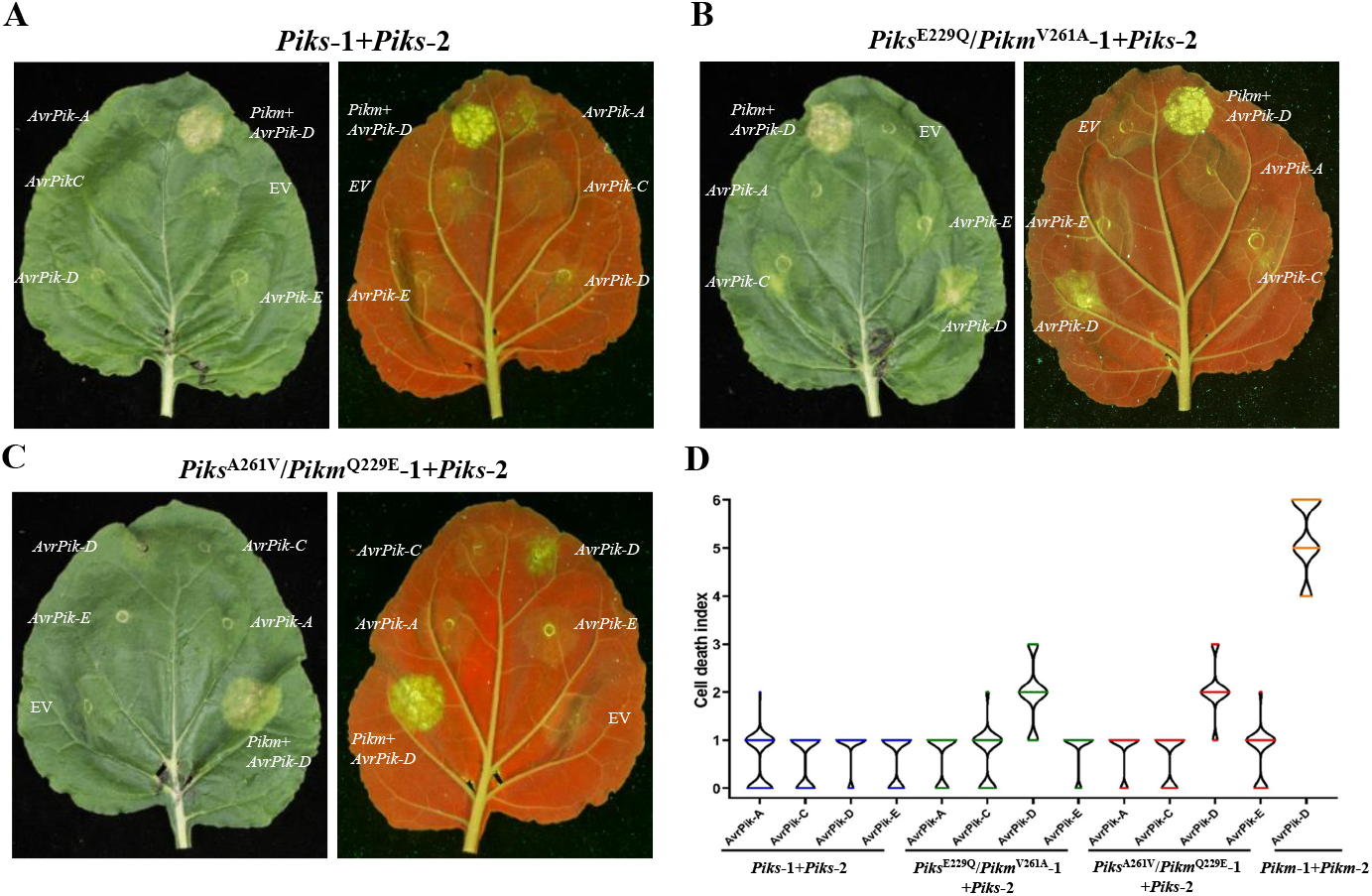
The *Piks* and *Piks* single amino acid substitution mutants mediated cell death response to *AvrPik* effector variants in *N. benthamiana*. (A-C), Representative leaf image showing *Piks, Piks*^E229Q^/*Pikm*^V261A^ and *Piks*^A261V^/*Pikm*^Q229E^ mediated cell death to *AvrPik* variants; Images showing autofluorescence are horizontally flipped to present the same leaf orientation as white light images; *Pikm*-mediated cell death with *AvrPik*-D is included as a positive control. (D) Box plots showing repeats of the cell death assay. For each sample, the number of repeats was 90. The centre line represents the median, the box limits are the upper and lower quartiles, the whiskers are the 1.5× interquartile range and all of the data points are represented as dots. The cell death scoring scale used is shown in Maqbool *et al*., 2015.

## DISCUSSION

In this study, we report the characterization of the rice NLR immune receptor *Piks* and investigate its ability to recognise *AvrPik* effector variants. An extensive genetic analysis using a set of authentic isogenic rice lines and rice blast isolates demonstrates that *Piks* is a novel functional allele conferring a distinct resistance spectrum from other allelic variants at the *Pik* locus. As is well established for other *Pik* genes, *Piks-1* and *Piks-2* are necessary and sufficient for *Piks*-mediated resistance. However, *Piks*, unlike its alleles, does not recognise any known *AvrPik* variants in rice. Instead, our results suggest it recognizes a novel *Avr* gene which is most likely dissimilar to *AvrPik* at the sequence and/or structural level. We performed reciprocal exchange of the two amino acid residues that differ between *Piks* and *Pikm* (at positions 229 and 261), which are both contained within the HMA domain, and tested their roles in transgenic rice towards sustaining distinct specificities of *Piks* and *Pikm*. In addition to genetic characterization, the physical binding activity of *Piks*-HMA and *Pikm*-mimicking mutants towards different *AvrPik* variants were investigated in both Y2H and analytical gel filtration assays. Finally, cell death assay between *Piks* mutants and *AvrPik* variants was conducted in *N. benthamiana*. These results reveal a complex scenario of interaction outcomes between *Piks* mutants and *AvrPik* variants using genetic and biochemical approaches.

### *Piks* confers a novel recognition specificity diverged from the canonical *Pik* NLRs

Clustering of multiple alleles at the same *R*-gene locus is evident in many crop species, such as the *L5*/*L6*/*L7* locus in flax (Dodds et al. 2004), the *MLA*1/7/9/10 locus in barley (Lu et al. 2016), and the *Pi2*/*Piz-t*/*Pi9*/*Pi50* locus in rice (Zhou et al. 2006; Qu et al. 2006; Su et al. 2015). Notably, these allelic variants frequently share close sequence similarity. For instance, the *MLA* and *Pi2/Piz-t/Pi9* alleles share over 90% and 96% sequence identity, respectively (Bauer et al. 2021; Zhou et al. 2007). Further, sequence variations within *R*-gene alleles leads to differential recognition cognate *Avr* genes with or without significant sequence similarity. For example, *Pi9* and *Piz-t* activate rice immunity against distinct sets of rice blast isolates by recognizing sequence unrelated *AvrPi9* and *AvrPiz-t*, respectively (Wu et al. 2015; Li et al. 2009). By contrast, *AvrL567-A* and *AvrL567-D*, recognized by *L5* and *L6*, respectively, share 92% amino acid sequence identity (Dodds et al. 2004, 2006). Different alleles of the rice *R*-gene *Pik* recognize at least one of five *AvrPik* variants, i.e., *AvrPik*-A to -E through so-called stepwise arms race co-evolution (Wu et al. 2014; Wang et al. 2017). *Piks* has been placed in the most ancestral position in the chronological order, without its functional verification at the molecular level (Wu et al. 2014). Here, we show that *Piks* shares characteristics of classical *Pik* alleles with respect to its genomic organization and sequence relatedness. However, unlike other *Pik* alleles, *Piks* does not recognize known *AvrPik* variants. Instead, it recognizes a novel *Avr* gene, which is presumably sequence unrelated to *AvrPik*. The finding that *Piks* confers an *AvrPik*-independent resistance specificity demonstrates a new interaction mode in addition to those previously shown between *Pik* alleles and *AvrPik* variants. This suggests that *Pik* variants have undergone distinct evolutionary mechanisms to recognize both sequences related and unrelated *Avr* effectors to activate immunity to specific pathotypes of the rice blast pathogen.

Our findings align with similar scenario illustrated in several *R*-gene loci which harbors multiple allelic variants, such as flax *L* locus (Dodds et al. 2004, 2006), barley *MLA* locus (Lu et al. 2016; Saur et al. 2019), Arabidopsis *RPP13* locus (Hall et al. 2009). In these *R*-gene loci, some allelic *R*-gene variants detect allelic *Avr* proteins, whereas others detect sequence unrelated *Avr* proteins, suggesting dynamic co-evolutionary mechanisms. Intriguingly, examples of sequence divergent but structurally related *Avr* proteins recognized by allelic *R*-gene variants have recently been reported. For example, five *AvrPm3* variants, specifically recognized by different allelic *PM3*, belong to a large group of proteins with low sequence homology but predicted structural similarities (Bourras et al. 2019). Further, protein structure analysis predicted that known *AVR*_*A*_ proteins specifically recognized by different allelic *MLA* variants each adopt RNase-like folds although they are sequence unrelated from one another (Bauer et al. 2021). Although the putative effector recognized by *Piks* is yet to be reported, and maybe sequence diverse from *AvrPik*, it is interesting to speculate they maybe structurally related, especially as many rice blast effectors whose structures have been determined (including all those that interact with host HMA proteins) adopt the MAX effector fold (De Guillen et al. 2015).

### Perturbation of HMA interfaces results in alteration of binding activity and strength to *AvrPik* proteins

The HMA domain of *Pik* alleles is the key determinant of specifically for recognizing *AvrPik* variants through direct binding (Maqbool et al. 2015; De la Concepcion et al. 2018, 2021). There is an overall correlation between recognition specificity and binding of Pik-HMA domains by AvrPik variants (Kanzaki et al. 2012; De la Concepcion et al. 2018). Structure analyses of the complexes of Pik-HMAs and AvrPik effectors identified three interfaces in the HMA domain, dominate the interaction with AvrPik effectors (De La Concepcion et al. 2018). Pikp mainly employs interface 2 to determine its binding with AvrPik-D, whereas Pikm mediates binding through robust interaction at interfaces 2 and 3 (De La Concepcion et al. 2018). The two amino acid residues at position 229 and 261 that distinguish Piks-1 from *Pikm-1* are positioned within interfaces 2 and 3, respectively, suggesting the likely importance of these two interfaces in the binding of Piks to AvrPik variants. Sequence alignment reveals that Glu229 and Ala261 (as found in Piks) are conserved in Pik-1-Ku/Ka and Pikp-1/Pi7-1/Pikh-1, respectively (Meng et al. 2021), implying that replacement of either Glu229 or Ala261 in *Piks*-1 might restore binding activity between derivative HMA and AvrPik proteins. Indeed, both the mutations Piks-1^E229Q^ and Piks-1^A261V^ supported physical binding to the AvrPik-D variant as measured by Y2H and analytical gel filtration with purified proteins. But this restoration of effector binding did not extend to the variants AvrPik-E nor -A (that bind Pikm). In a recent study, *Pikg*, which originates from a wild rice species and differs from *Pike* by a single amino acid substitution (D229E) in the HMA domain, controls a distinct resistance spectrum (Meng et al. 2021). However, sequence changes at these equivalent positions in the HMA domain may not be a necessary result in the functional divergence. For example, *Pikm*-1 and *Pi1*-5 confer indistinguishable specificities although they differ from each other by a single amino acid change (Gln229Asp) at the equivalent position (Figure S5).

### Varied outcomes of interaction between *Piks* alleles and *AvrPik* variants

In this study, we used three different approaches to characterize the function of *Piks* and the mutants *Piks*-1^E229Q^/*Pikm*-1^V261A^ and *Piks*-1^A261V^/*Pikm*-1^Q229E^, protein-protein binding assay via Y2H and gel filtration analyses, cell death assays in *N. benthamiana*, and genetic complementation tests in rice (Table S3). The results reveal an overall consistent outcome derived from functional characterization using different approaches. Piks-1 shows no binding activity with tested AvrPik proteins in either Y2H or analytical gel filtration. It does not induce cell death in *N. benthamiana* when co-expressed with different *AvrPik* effectors, and cannot complement resistance in rice to isolates containing different *AvrPik* effectors (Table S3). Likewise, a consistent result is observed for the functional characterization of *Piks*-1^E229Q^/*Pikm*-1^V261A^. However, Piks-1^A261V^/Pikm-1^Q229E^ shows a gain of binding activity to the effector variant AvrPik-D and induces cell death when co-expressed with *AvrPik*-D in *N. benthamiana*, but it could not activate resistance against blast isolates containing *AvrPik*-*D*. The possibility that *Piks*-1^A261V^/*Pikm*-1^Q229E^ is not expressed properly is further excluded since the same transgenic lines retain the resistance against V86010. Therefore, we conclude that *Piks*-1^A261V^/*Pikm*-1^Q229E^ is incapable of activating immunity against rice blast isolates with *AvrPik*-D although it appears functional in both protein-protein binding and ectopic cell death assays. It is still elusive whether this similar observation could be applicable to other *R*-*Avr* functional studies. However, it provides a specific case illustrating where ectopic expression assays might blur the true function of particular *R* genes in recognition and immunity.

## MATERIALS AND METHODS

### Plant materials and growth condition

The F_2_ population derived from a cross between the *Piks* gene monogenic line IRBLKs-F5 and its recipient rice variety Lijiangxintuanheigu (LTH) were used for mapping and co-segregation analysis. The rice cultivar Nipponbare was used as a host cultivar for gene transformation. The *Pik* allelic gene monogenic lines in LTH background IRBK1-CL (*Pi1*), IRBL7-M (*Pi7*), IRBLK-Ka (*Pik*), IRBLKm-Ts (*Pikm*), and IRBLKp-K60 (*Pikp*) were used to evaluate the resistance specificity.

Rice seeds were surface-sterilized with sodium hypochlorite and plated on petri dishes covered with wet filter paper. Seeds were germinated at 37°C in a growth chamber with a 12-h/12-h light/dark cycle. Standard growth on soil in a growth chamber was under a 16h/8h light/dark cycle with 30°C/22°C, if not otherwise indicated.

### *M.oryzae* materials and pathogenicity assay

The field *M. oryzae* isolates JMB8401, 5008-3, 9482-1-1, CA89 and V86010 were used in this study. To create isogenic lines of *M. oryzae* harboring different *AvrPik* alleles, we transformed R88-002 individually with expression vectors for *AvrPik*-A, *AvrPik*-B, *AvrPik*-C, *AvrPik*-D, *AvrPik*-E and *AvrPik*-F. We amplify the native promoter (∼1.5kb) and ORF (∼0.3kb) of *AvrPik*-A to -F from the *M. oryzae* isolates 98-06, Guy 11, V86010 and R01-1, and digested using *SalI* and *EcoRI* and connected to pCB1532-attR vector, individually.

The *AvrPik* gene replacement constructs pMD19-T-*AvrPik*-KO was generated according to the homologous recombination principle as described previously (Zhang et al. 2010). The spacers for Cas9-gRNA vectors were designed with the gRNA designer program for best on-target scores (http://grna.ctegd.uga.edu/). The sense and antisense oligonucleotides of the selected gRNA spacers were synthesized and annealed to generate the corresponding gRNA spacers, and cloned between the two *BsmBI* sites of pCas9-tRp-gRNA by Golden Gate cloning (NEB) (Arazoe et al. 2015) to generate the pCas9-tRpgRNA-*AvrPik* constructs. Then, the gene replacement vector pMD19-T-*AvrPik*-KO and its corresponding pCas9-tRpgRNA-*AvrPik* vector were co-transformed into protoplasts of V86010 to obtain the *AvrPik* gene deletion mutants. Hygromycin-resistant transformants were screened by PCR and selected for phenotype analysis. Primers used in this section are listed in Table S2.

For fungal transformation, protoplasts were isolated and transformed with the PEG method as described previously (Sweigard et al. 1995). Media were supplemented with chlorimuron-ethyl/hygromycin B at 300 µg/ml in order to select chlorimuron-ethyl/ hygromycin resistant transformants. All the resultant transformants were subjected to either PCR validation using specific primers (Table S2) or Southern hybridization to ascertain successful single-copy integration.

Rice seedlings at the 3-4 leaf stage were used for spray inoculation. The spore suspension with 10^5^ spores/ml was applied to plants using an airbrush connected to a source of compressed air. After inoculation, plants were held in the dark room for 24 h with 95–100% relative humidity and 24°C. Then, plants were transferred to a greenhouse where the temperature was maintained at 25–28 °C and humidity was keep at 70–90%. Six to Seven days after inoculation, disease symptoms were evaluated using a standard 0–5 scale (IRRI 2002).

### DNA extraction, PCR primers and sequencing analysis

Genomic DNA of all the plant materials was extracted from fresh leaves using CTAB method. For fungal DNA extractions, mycelia were harvested from 3-day-old cultures grown in liquid CM at 28°C with shaking at 150 rpm. General molecular biology techniques for nucleic acid analysis were performed according to standard protocols. We used PCR primer pair RGA4-F3/RGA4-R3 to genotype the F_2_ population for co-segregation analysis. The primer pairs Piks-1-F/Piks-1-R and Piks-2-F/ Piks-2-R were used to define the candidate genome region of *Piks*, to check the presence of transgenes in transgenic lines (Table S2). For *AvrPik* gene haplotype analysis in wild *M. oryzae* isolates, the primer AvrPik-F/AvrPik-R was use for PCR amplification, and selected PCR products were subjected to Sanger sequencing. The sequence of the primers were list at Table S1.

### Candidate gene cloning and complementation analysis

As reported previously, the two genes of *Pik* (*Pik*-1 and *Pik*-2) alleles working together to trigger *Pik* mediated resistance. From the genomic DNA of IRBLKs-F5, we used a high-fidelity Taq (Phanta Super-Fidelity DNA Plomerase, Vazyme) to amplify a 10.5-kb fragement, which contain the whole *Piks*-1 coding sequence, 5’-untranslated region (UTR) and 3’-UTR. The whole *Piks*-2 coding sequence, 5’-untranslated region (UTR) and 3’-UTR, which was 8.9-kb were also amplified using the same method. The two amplified products were individually inserted into the AscI and PacI site of the vector pCambia 1305.2 to form constructs for transformation. These constructs were validated by comparison of their insert sequences with the sequence published at NCBI (HQ662329). Mutagenesis of *Piks*-1 was performed using the Mut Express MultiS Fast Mutagenesis Kit V2 (VAZYME, Nanjing, China). The two mutant *Piks*-1^229Q^/*Pikm*-1^261A^ and *Piks*-1^261V^/*Pikm*-1^229E^ were confirmed successful gene mutation by select the plasmids for sequencing.

The four constructs each containing the *Piks*-1, *Piks*-1^E229Q^/*Pikm*-1^V261A^, *Piks*-1^A261V^/*Pikm*-1^Q229E^ and *Piks*-2, individually, were transferred into Nipponbare callus by Agrobacterium-mediated transformation. The Primary transgenic plants (T0 plants) regenerated from hygromycin-resistant calluses were grown in an isolated greenhouse. The T0 plants of *Piks*-1, *Piks*-1^E229Q^/*Pikm*-1^V261A^ and *Piks*-1^A261V^/*Pikm*-1^Q229E^ were checked by PCR assay with the *Piks*-1 specific primer pair Piks-1-F/Piks-1-R and the *Piks*-2 T0 plants were checked by specific primer pair Piks-2-F/ Piks-2-R (Table S2).

### Gene cloning: heterologous protein production, Y2H and in planta expression

For in vitro studies, *Piks*-HMA, *Piks*-1^229Q^/*Pikm*-1^261A^-HMA and *Piks*-1^261V^/*Pikm*-1^229E^ -HMA (residues Gly186 to Asp264, codon optimized for expression in *E. coli*) was generated by introducing the Glu229Gln and Ala261Val in *Pikm*-HMA by site-directed mutagenesis, and followed by cloning into the vector pOPIN-M (Maqbool et al. 2015; De la Concepcion et al. 2018). The expression constructs of *Pikm*-HMA and *AvrPik* used in this study were described previously (Maqbool et al. 2015; De la Concepcion et al. 2018).

For yeast-2-hybrid analyses, the *Pikm*-HMA, *Piks*-HMA, *Piks*-1^E229Q^/*Pikm*-1^V261A^-HMA and *Piks*-1^A261V^/*Pikm*-1^Q229E^-HMA (residues Gly186 to Ile300) was synthesized and supplied in the pGBKT7, the *AvrPik*-A, -C, -D and -E (residues Glu22 to Phe113, lacking the signal peptide) was synthesized (Sangon Biotech) and supplied in the pGADT7.

For protein expression in planta, the *Pikm*-HMA, *Piks*-HMA, *Piks-1*^E229Q^/*Pikm-1*^V261A^-HMA and *Piks-1*^A261V^/*Pikm-1*^Q229E^-HMA domain was assembled into a full-length NLR construct using Golden Gate cloning (Maqbool et al. 2015; De la Concepcion et al. 2018) and then into the plasmid pICH47742 with a C-terminal 6xHis/3xFLAG tag. The expression was driven by the A. tumefaciens Mas promoter and terminator.

### Expression and purification of proteins for in vitro binding studies

The *Pikm*-HMA, *Piks*-HMA, *Piks-1*^E229Q^/*Pikm-1*^V261A^-HMA and *Piks-1*^A261V^/*Pikm-1*^Q229E^-HMA with the 6xHis-MBP-tag were produced in *E. coli* SHuffle cells using the protocol described previously (Maqbool et al. 2015; De la Concepcion et al. 2018). Cell cultures were incubated in auto induction media at 30°C for 5-7 hours and then at 16°C overnight. Cells were harvested by centrifugation and resuspended in the medium used previously (Maqbool et al. 2015; De la Concepcion et al. 2018). The suspended Cells were sonicated and then centrifugated at 40000 x *g* for 30min, the supernatant lysate was applied to a Ni^2+^-NTA column connected to an AKTA Xpress purification system (GE Healthcare) (Maqbool et al. 2015; De la Concepcion et al. 2018). Proteins were step-eluted with elution buffer and directly injected onto a Superdex 75 26/600 gel filtration column pre-equilibrated 20mM HEPES pH 7.5, 150 mM NaCl. Purification tags were then removed by incubating with 3C protease (10 μg/mg fusion protein) overnight at 4°C followed by passing through tandem Ni^2+^-NTA and MBP Trap HP columns (GE Healthcare). The flow-through was concentrated as appropriate and loaded on a Superdex 75 26/600 gel filtration column for final purification and buffer exchange into 20 mM HEPES pH 7.5, 150 mM NaCl (Maqbool et al. 2015; De la Concepcion et al. 2018).

The *AvrPik* effectors, with a 3C protease-cleavable N-terminal SUMO tag and a non-cleavable C-terminal 6xHis tag, were produced in and purified from *E. coli* SHuffle cells as previously described (Maqbool et al. 2015; De la Concepcion et al. 2018). All protein concentrations were determined using a Direct Detect Infrared Spectrometer (Merck).

### Analytical gel filtration

Analytical size exclusion chromatography was performed at 4°C using a Superdex 75 10/300 gel filtration column (GE Healthcare) pre-equilibrated in 20 mM HEPES pH 7.5 and 150 mM NaCl (Maqbool et al. 2015; De la Concepcion et al. 2018). Samples were centrifuged before loading. A 100 μl of the sample was injected at a flow rate of 0.8 ml/min and 0.5 ml fractions were collected for analysis by SDS-PAGE gels. Proteins were mixed and incubated on ice for 60 min before loading for study complex formation (Maqbool et al. 2015; De la Concepcion et al. 2018).

### Yeast two-hybrid assay

We use the Matchmaker Gold Yeast Two-Hybrid System (Takara Bio USA) to detect protein–protein interactions between *Pik*-HMAs and *AvrPik* effectors. The DNA encoding the *Piks*-HMAs in pGBKT7 was co-transformed with either the individual *AvrPik* variants in pGADT7 into chemically competent Saccharomyces cerevisiae Y2HGold cells (Takara Bio USA). The single colonies grown on selection plates were inoculated in SD^-Leu-Trp^ plate and grown at 30 °C for overnight. Saturated culture was then used to make serial dilutions of optical density at 600 nm (OD600) 1, 1^-1^, 1^-2^ and 1^-3^, respectively. Five microlitre of each dilution was spotted on a SD^-Leu-Trp^ plate as a growth control and also on a SD^-Leu-Trp-Ade-His^ plate containing X-α-gal and aureobasidine, as detailed in the user manual. After incubated at 30 °C for 60–72 h, the plates were imaged. Each experiment was repeated three times, with similar results.

### *N. benthamiana* cell death assays

For agroinfiltration in *N. benthamiana*, Agrobacterium tumefaciens GV3101 was transformed with the relevant T-DNA constructs. Leaves of 4-week-old *N. benthamiana* plants grown at 22–25 °C with high light intensity were agroinfiltrated using a needleless syringe (Maqbool et al. 2015; De la Concepcion et al. 2018). *Piks*-1(*Piks*-1, *Piks*-1^E229Q^/*Pikm*-1^V261A^-1 and *Piks*-1^A261V^/*Pikm*-1^Q229E^-1), *Piks*-2, *AvrPik* and P19 were mixed at OD600 0.4, 0.4, 0.6 and 0.1, respectively (Maqbool et al. 2015; De la Concepcion et al. 2018). Detached leaves were imaged at 5 dpi from the adaxial side of the leaves for white light image and abaxial side of the leaves for UV images. Pictures are representative of three independent experiments, with internal repeats. The cell death index used for scoring is as presented previously (Maqbool et al. 2015; De la Concepcion et al. 2018). The scoring for all replicas is presented as box plots, which were generated using R v3.4.3 (https://www.r-project.org/) and the GraphPad Prism 9.0 (https://www.graphpad-prism.cn/). The centre line represents the median, the box limits are the upper and lower quartiles, the whiskers are the 1.5× interquartile range and all of the data points are represented as dots (Maqbool et al. 2015; De la Concepcion et al. 2018).

## ACKNOWLEDGMENTS

This work was supported, in part, by grants from the National Natural Science Foundation of China for J.W. (U20A2021), Major Science and Technology Project of Hunan Province (2021NK1001), and Talent promotion project of Hunan Association for Science and Technology (2019TJ-N03).

## AUTHOR CONTRIBUTIONS

G.X., J.W. and B.Z. conceived and designed the experiments. G.X., W.W., M.L., Y.L., carried out the experiments. G.X., J.L., M.F. and Z.Y. analyzed the data. X.Z., Z.Z. and G.L. provided technical assistance. G.X., M.B., J.W. and BZ wrote the manuscript. All authors read and approved the final manuscript.

## SUPPORTING INFORMATION

**Figure S1**. Detection of the rice blast strain R88-002 transformed with six *AvrPik* variants (*AvrPik*-A to –F) by PCR assay.

**Figure S2**. Gene specific marker which can distinguishes *Piks* and *Pikm* from LTH (Lijiangxintuanheigu) and other five *Pik* alleles. Lane 1, IRBLks-F5 (*Piks*); Lane 2, LTH; Lane 3, IRBLkm-TS (*Pikm*); Lane 4 IRBLkh-K3 (*Pikh*); Lane 5, IRBLkp-K60 (*Pikp*); Lane 6, IRBLK-Ka (*Pik*); Lane 7, IRBL1-CL (*Pi1*); Lane 8, IRBL7-M (*Pi7*); L, DNA Ladder.

**Figure S3**. The rice blast isolate V86010 contain both *AvrPik*-D and *AvrPik*-E. (A) Sequence alignment of *AvrPik* in V86010, *AvrPik*-D and *AvrPik*-E. (B) Sequencing results of *AvrPik* PCR amplicon. The red triangle marked out the location of sequence difference and double peak of the sequencing results.

**Figure S4**. PCR amplification results of the three *AvrPik* knock out mutant strains of V86010. AvrPik-In-F/R primer pair cover the coding region of *AvrPik* while the primer pair AvrPik-out-F/Hoh-R amplified the replacement fragment and promoter of the *AvrPik*.

**Figure S5**. Alignment of amino acid sequence encoded by *Pik*-1 alleles.

**Figure S6**. Amino acid residue polymorphisms among *Pik*-2 alleles. *Dots* represent residues identical to those in *Piks*-2.

**Figure S7**. Representative SDS-PAGE gels of relevant fractions. (A) SDS-PAGE gel of *AvrPik*-D protein. (B) SDS-PAGE gel of *Pikm* and *AvrPik*-D protein complex.

**Table S1**. Pathogenicity of different *M. oryzae* isolates toward *Pik* allelic gene monogenic lines in Lijiangxintuanheigu (LTH).

**Table S2**. Primers used in this study.

**Table S3**. The various interactions and phenotypes between *Piks-1, Piks* mutants and *AvrPik* variants in this study.

