## supplemental information for "The *Piks* allele of the NLR immune receptor *Pik* breaks the recognition of *AvrPik* effectors of the rice blast fungus"

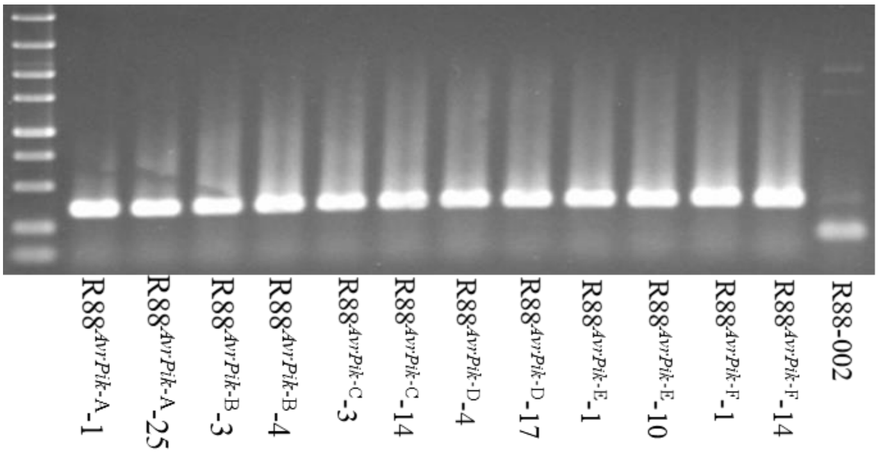


**Figure S1.** Detection of the rice blast strain R88-002 transformed with six *AvrPik* variants (*AvrPik*-A to –F) by PCR assay.

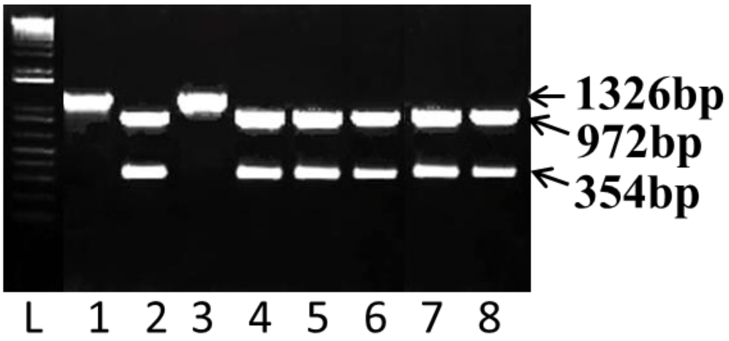


**Figure S2.** Gene specific marker which can distinguishes *Piks* and *Pikm* from LTH (Lijiangxintuanheigu) and other five *Pik* alleles. Lane 1, IRBLks-F5 (*Piks*); Lane 2, LTH; Lane 3, IRBLkm-TS (*Pikm*); Lane 4 IRBLkh-K3 (*Pikh*); Lane 5, IRBLkp-K60 (*Pikp*); Lane 6, IRBLK-Ka (*Pik*); Lane 7, IRBL1-CL (*Pi1*); Lane 8, IRBL7-M (*Pi7*); L, DNA Ladder.

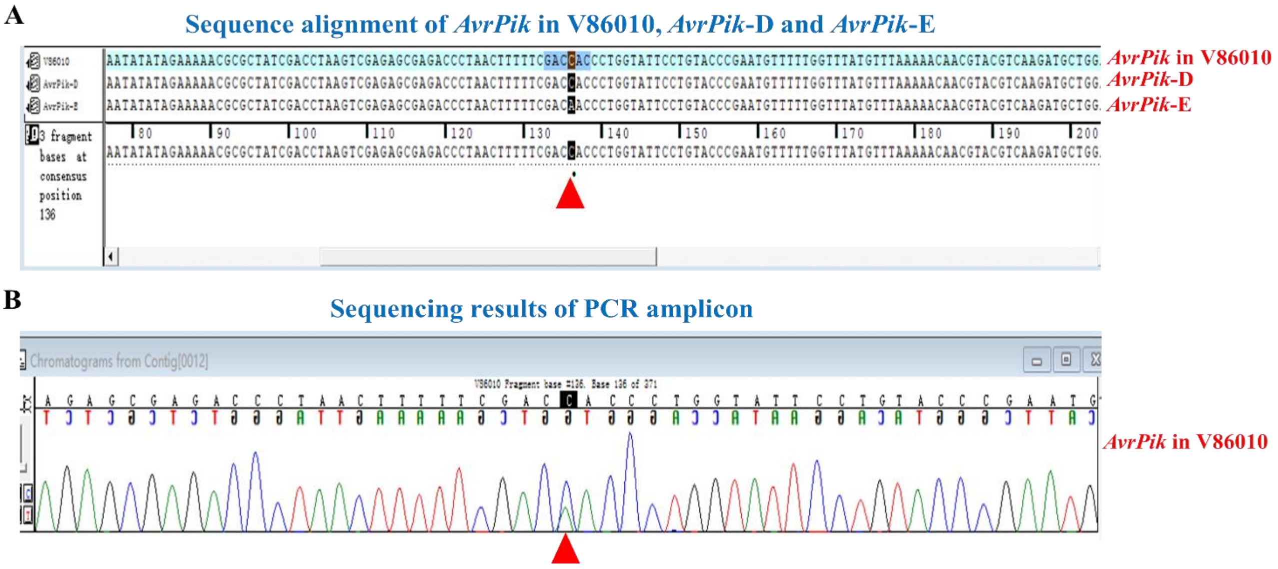


**Figure S3.** The rice blast isolate V86010 contain both *AvrPik*-D and *AvrPik*-E. (A) Sequence alignment of *AvrPik* in V86010, *AvrPik*-D and *AvrPik*-E. (B) Sequencing results of *AvrPik* PCR amplicon. The red triangle marked out the location of sequence difference and double peak of the sequencing results.

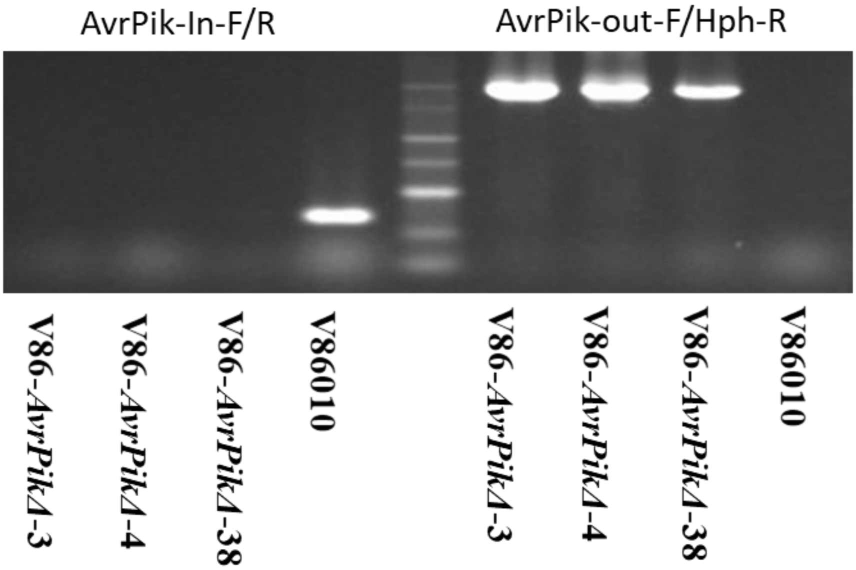


**Figure S4.** PCR amplification results of the three *AvrPik* knock out mutant strains of V86010. AvrPik-In-F/R primer pair cover the coding region of *AvrPik* while the primer pair AvrPik-out-F/Hoh-R amplified the replacement fragment and promoter of the *AvrPik*.

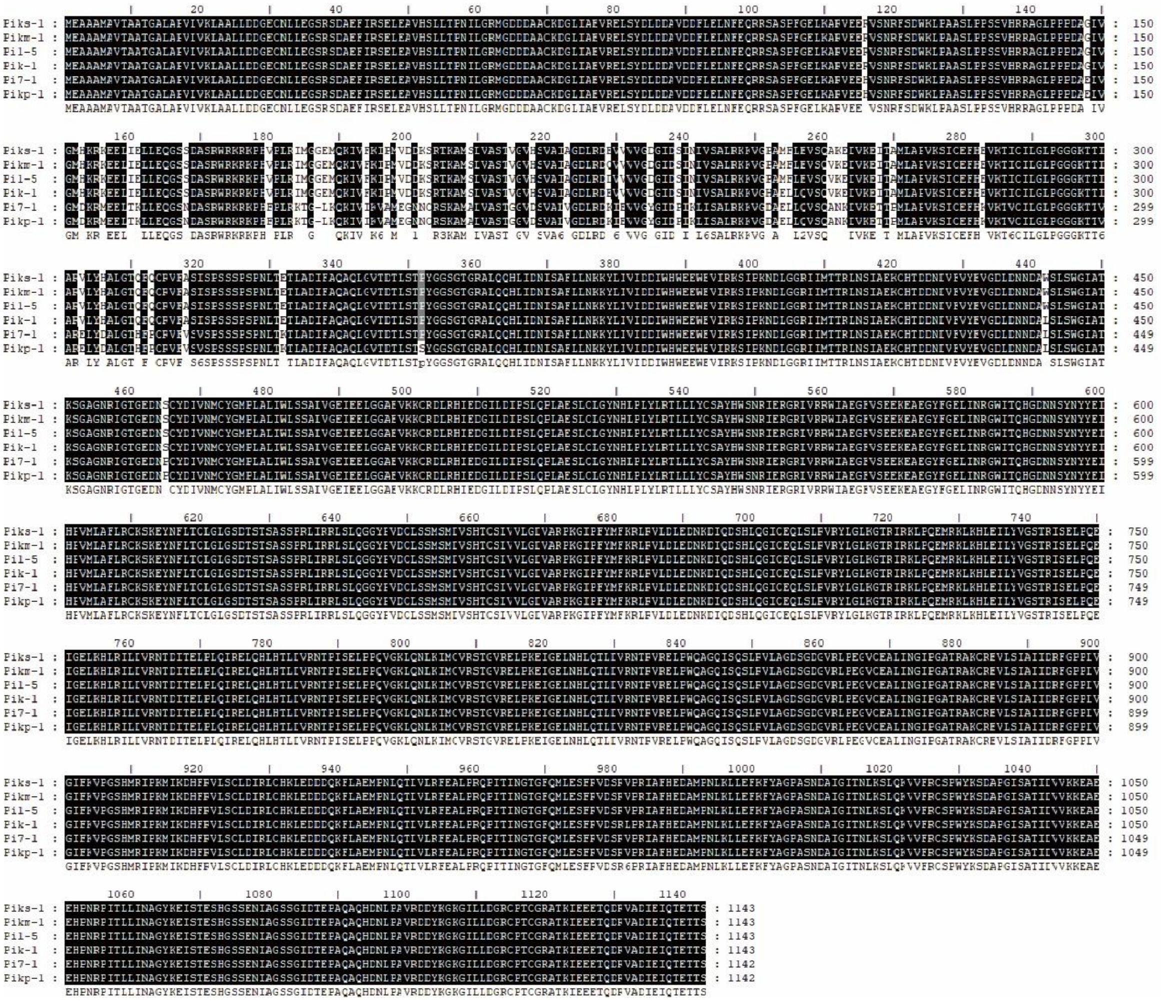

**Figure S5.** Alignment of amino acid sequence encoded by *Pik-1* alleles.

| | NBS | | LRR | |
|---|---|---|---|---|
| | 230 | 434 | 627 | 754 |
| ***Piks-2*** | **E** | **S** | **V** | **E** |
| ***Pikm-2*** | • | • | • | • |
| ***Pik-2*** | • | • | • | • |
| ***Pi1-6*** | • | • | • | **V** |
| ***Pikp-2*** | **D** | **T** | **M** | • |
| ***Pi7-2*** | **D** | **T** | **M** | • |

**Figure S6.** Amino acid residue polymorphisms among *Pik-2* alleles. *Dots* represent residues identical to those in *Piks-2*.

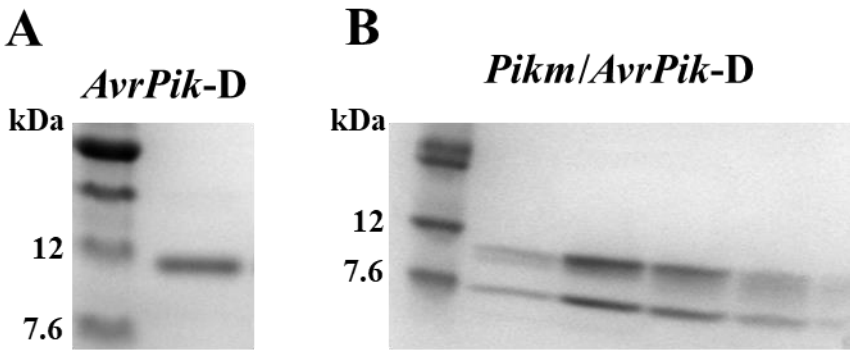


**Figure S7.** Representative SDS-PAGE gels of relevant fractions. (A) SDS-PAGE gel of *AvrPik*-D protein. (B) SDS-PAGE gel of *Pikm* and *AvrPik*-D protein complex.

Table S1. Pathogenicity of different *M. oryzae* isolates toward *Pik* allelic gene monogenic lines in Lijiangxintuanheigu (LTH)

| Seq | Isolate | IRBLks-F5 *Piks* | IRBLk-ka *Pik* | IRBLkp-K60 *Pikp* | IRBL1-CL *Pi1* | IRBL7-M *Pi7* | IRBLkm-Ts *Pikm* | LTH |
|---|---|---|---|---|---|---|---|---|
| 1 | PO6-6 | S | R | R | R | R | R | S |
| 2 | B90018 | S | R | R | R | R | R | S |
| 3 | B90002 | S | R | R | R | R | R | S |
| 4 | Ca89 | S | R | R | R | R | R | S |
| 5 | IK81-3 | S | R | R | R | R | R | S |
| 6 | Ca41 | S | R | R | R | R | R | S |
| 7 | M64-1-3-9-1 | S | S | S | S | S | S | S |
| 8 | V86010 | R | R | R | R | R | R | S |
| 9 | C9228-37 | S | R | R | R | R | R | S |
| 10 | V86046 | S | R | R | R | R | R | S |
| 11 | M39-1-3-8-1 | S | S | S | S | S | S | S |
| 12 | IK81-25 | S | S | S | S | S | S | S |
| 13 | 92318-4 | S | S | S | S | S | S | S |
| 14 | M64-7-2-8-2 | S | R | R | R | R | R | S |
| 15 | JMB840106 | S | S | R | R | S | R | S |
| 16 | B90038 | S | R | R | R | S | R | S |
| 17 | C923-49 | S | R | R | R | R | R | S |
| 18 | JMB840197 | S | R | R | R | R | R | S |
| 19 | JMB8401 | S | S | S | S | S | S | S |
| 20 | JMB840610 | S | S | S | S | S | S | S |
| 21 | Ca91 | S | R | S | R | S | R | S |
| 22 | PO83-Z1-30 | S | S | S | S | S | S | S |
| 23 | 9239-4 | S | S | S | S | S | S | S |
| 24 | M101-1-2-9-1 | S | S | S | S | S | S | S |
| 25 | M36-1-3-10-1 | S | S | S | S | S | S | S |
| 26 | BN111 | S | R | R | R | R | R | S |
| 27 | B90103 | S | R | R | S | R | S | S |
| 28 | 92329-9 | S | S | S | S | S | S | S |
| 29 | JMB840298 | S | S | S | S | S | S | S |
| 30 | Pi9-1 | S | S | S | S | S | S | S |
| 31 | BN209 | S | S | S | S | S | S | S |
| 32 | Pi9-4 | S | S | S | S | S | S | S |
| 33 | B90039 | S | R | R | R | R | R | S |
| 34 | B90020 | S | R | R | R | R | R | S |
| 35 | B90298 | S | R | R | R | R | R | S |
| 36 | B90321 | S | R | R | R | R | R | S |
| 37 | B90075 | S | R | R | R | R | R | S |
| 38 | B90033 | S | R | R | R | R | R | S |
| 39 | 9237-36 | S | R | R | R | R | R | S |

| | | | | | | | | |
|---|---|---|---|---|---|---|---|---|
| 40 | 92319-9 | S | R | R | R | R | R | S |
| 41 | B90036 | S | R | R | R | R | R | S |
| 42 | CBN214-1 | S | R | R | R | R | R | S |
| 43 | V850229 | S | R | R | R | R | R | S |
| 44 | C9210-4 | S | R | R | R | R | R | S |
| 45 | JMB840495 | S | R | S | R | R | R | S |
| 46 | C9239-4 | S | R | R | R | R | R | S |
| 47 | M101-7-2-1-1 | S | S | S | R | R | R | S |
| 48 | 9248-6 | S | S | R | R | R | R | S |
| 49 | JMB840112 | S | R | R | R | R | R | S |
| 50 | 4029-1 | S | S | R | S | S | S | S |
| 51 | 5008-3 | S | R | R | R | R | R | S |
| 52 | 5029-1 | S | S | R | S | S | S | S |
| 53 | 5033-1 | S | S | R | S | S | S | S |
| 54 | 5047-3 | S | S | R | S | S | S | S |
| 55 | 5054-3 | S | S | R | S | S | S | S |
| 56 | 5092-3 | S | R | R | R | R | R | S |
| 57 | 5093-3 | S | S | R | S | S | S | S |
| 58 | 5127-1 | S | R | R | R | R | R | S |
| 59 | 5131-2 | S | R | R | R | R | R | S |
| 60 | 5140-2 | S | S | R | S | S | S | S |
| 61 | 5156-3 | S | S | R | S | S | S | S |
| 62 | 5167-1 | S | R | R | R | R | R | S |
| 63 | 6003-3 | S | R | R | R | R | R | S |
| 64 | 6006-1 | S | R | R | R | R | R | S |
| 65 | 6016-2 | S | R | R | R | R | R | S |
| 66 | 6021-2 | S | S | R | S | S | S | S |
| 67 | 6030-3 | S | S | R | S | S | S | S |
| 68 | 6033-3 | S | S | R | S | S | S | S |
| 69 | 6050-3 | S | R | R | R | R | R | S |
| 70 | 6054-1 | S | S | R | S | S | S | S |
| 71 | 6057-3 | S | S | R | S | S | S | S |
| 72 | 6065-2 | S | S | R | S | S | S | S |
| 73 | 6085-3 | S | S | R | S | S | S | S |
| 74 | 6116-2 | S | S | R | S | S | S | S |
| 75 | 6119-1 | S | S | R | S | S | S | S |
| 76 | 6161-1 | S | R | R | R | R | R | S |
| 77 | 7024-1 | S | S | R | S | S | S | S |
| 78 | 7068-2 | S | S | R | R | S | R | S |
| 79 | 8036-2 | S | S | R | S | S | S | S |
| 80 | 8081-2 | S | S | R | S | S | S | S |
| 81 | 9048-1 | S | S | R | S | S | S | S |
| 82 | 9056-1 | S | S | R | S | S | S | S |
| 83 | 9113-1 | S | S | R | S | S | S | S |

| | | | | | | | | |
|---|---|---|---|---|---|---|---|---|
| 84 | 9113-2 | S | S | R | S | S | S | S |
| 85 | 9121-1 | S | S | R | S | S | S | S |
| 86 | 9124-1 | S | S | R | S | S | S | S |
| 87 | 9126-1 | S | R | R | R | R | R | S |
| 88 | 9149-1A | S | S | R | S | S | S | S |
| 89 | 9157-3B | S | S | R | S | S | S | S |
| 90 | 9186-1 | S | S | R | S | S | S | S |
| 91 | 9190-3 | S | S | R | S | S | S | S |
| 92 | 9193-1A | S | S | R | S | S | S | S |
| 93 | 9202-1 | S | S | R | S | S | S | S |
| 94 | 9212-1 | S | S | R | S | S | S | S |
| 95 | 9221-3 | S | S | R | S | S | S | S |
| 96 | 9233-1 | S | S | R | S | S | S | S |
| 97 | 9244-3 | S | R | R | R | R | R | S |
| 98 | 9245-1 | S | S | R | S | S | S | S |
| 99 | 9250-2 | S | S | R | S | S | S | S |
| 100 | 9252-1 | S | S | R | S | S | S | S |
| 101 | 9257-1 | S | S | R | S | S | S | S |
| 102 | 9262-3 | S | S | R | S | S | S | S |
| 103 | 9264-1A | S | S | R | S | S | S | S |
| 104 | 9265-1 | S | S | R | S | S | S | S |
| 105 | 9277-3B | S | S | R | S | S | S | S |
| 106 | 9293-1 | S | S | R | S | S | S | S |
| 107 | 9304-2 | S | S | R | S | S | S | S |
| 108 | 9310-1 | S | S | R | S | S | S | S |
| 109 | 9316-1 | S | S | R | S | S | S | S |
| 110 | 9323-1 | S | S | R | S | S | S | S |
| 111 | 9324-1 | S | S | R | S | S | S | S |
| 112 | 9333-3 | S | S | R | S | S | S | S |
| 113 | 9334-3 | S | S | R | S | S | S | S |
| 114 | 9403-3 | S | S | R | S | S | S | S |
| 115 | 9406-3 | S | R | R | R | R | R | S |
| 116 | 9440-1 | S | S | R | S | S | S | S |
| 117 | 9459-3 | S | S | R | S | S | S | S |
| 118 | 9475-1-3 | S | R | R | R | R | R | S |
| 119 | 9482-1-3 | S | S | S | R | S | R | S |
| 120 | 9497-3 | S | R | R | R | R | R | S |
| 121 | 9533-1 | S | S | R | S | S | S | S |
| 122 | BZ64-1 | S | S | R | S | S | S | S |
| 123 | MO15-1 | S | R | R | R | R | R | S |
| 124 | MO15-2 | S | R | R | R | R | R | S |
| 125 | MO15-3 | S | R | R | R | R | R | S |
| 126 | MO15-4 | S | R | R | R | R | R | S |
| 127 | MO15-5 | S | S | R | R | S | R | S |

| | | | | | | | | |
|---|---|---|---|---|---|---|---|---|
| 128 | MO15-6 | S | S | S | S | S | S | S |
| 129 | MO15-7 | S | S | S | S | S | S | R |
| 130 | MO15-8 | S | S | R | R | S | R | S |
| 131 | MO15-9 | S | S | S | S | S | S | S |
| 132 | MO15-10 | S | S | R | R | S | R | S |
| 133 | MO15-11 | S | R | R | R | S | R | S |
| 134 | MO15-12 | S | S | R | R | S | R | S |
| 135 | MO15-13 | S | S | S | S | S | S | S |
| 136 | MO15-14 | S | R | R | R | R | R | S |
| 137 | MO15-15 | S | S | S | S | S | S | S |
| 138 | MO15-16 | S | S | R | R | S | R | S |
| 139 | MO15-17 | S | R | R | R | R | R | S |
| 140 | MO15-19 | S | S | S | S | S | S | S |
| 141 | MO15-20 | S | R | S | R | S | S | S |
| 142 | MO15-21 | S | R | R | R | R | R | S |
| 143 | MO15-22 | S | S | R | R | S | R | S |
| 144 | MO15-23 | S | S | S | S | S | S | S |
| 145 | MO15-24 | S | R | S | R | S | R | S |
| 146 | MO15-25 | S | R | R | R | R | R | S |
| 147 | MO15-26 | S | S | S | S | S | S | S |
| 148 | MO15-27 | S | S | S | S | S | S | S |
| 149 | MO15-28 | S | S | R | R | R | R | S |
| 150 | MO15-29 | S | S | R | R | S | R | S |
| 151 | MO15-30 | S | S | S | R | S | R | S |
| 152 | MO15-31 | S | R | R | R | R | R | S |
| 153 | MO15-32 | S | R | R | R | R | R | S |
| 154 | MO15-33 | S | S | R | R | S | R | S |
| 155 | MO15-34 | S | S | R | R | S | R | S |
| 156 | MO15-35 | S | R | R | R | R | R | S |
| 157 | MO15-36 | S | R | R | R | R | R | S |
| 158 | MO15-37 | S | S | R | R | S | R | S |
| 159 | MO15-38 | S | S | S | S | S | S | S |
| 160 | MO15-39 | S | R | R | R | R | R | S |
| 161 | MO15-40 | S | R | R | R | R | R | S |
| 162 | MO15-41 | S | S | R | R | S | R | S |
| 163 | MO15-42 | S | R | R | R | R | R | S |
| 164 | MO15-43 | S | R | R | R | R | R | S |
| 165 | MO15-44 | S | R | R | R | R | R | S |
| 166 | MO15-45 | S | S | S | S | S | S | S |
| 167 | MO15-46 | S | R | R | R | R | R | S |
| 168 | MO15-47 | S | S | R | R | S | S | S |
| 169 | MO15-48 | S | R | R | R | R | R | S |
| 170 | MO15-49 | S | S | R | R | S | R | S |
| 171 | MO15-50 | S | S | R | R | S | R | S |

| | | | | | | | | |
|---|---|---|---|---|---|---|---|---|
| 172 | MO15-51 | S | R | R | R | R | R | S |
| 173 | MO15-52 | S | S | S | S | S | S | S |
| 174 | MO15-53 | S | S | S | R | S | R | S |
| 175 | MO15-54 | S | R | R | R | R | R | S |
| 176 | MO15-55 | S | S | R | S | S | S | S |
| 177 | MO15-56 | S | R | R | R | R | R | S |
| 178 | MO15-56 | S | R | R | R | R | R | S |
| 179 | MO15-57 | S | R | R | R | R | R | S |
| 180 | MO15-58 | S | R | R | R | R | R | S |
| 181 | MO15-59 | S | S | R | R | S | R | S |
| 182 | MO15-60 | S | R | R | R | R | R | S |
| 183 | MO15-61 | S | S | R | R | S | R | S |
| 184 | MO15-62 | S | S | R | R | S | R | S |
| 185 | MO15-63 | S | S | R | R | S | R | S |
| 186 | MO15-64 | S | R | R | R | R | R | S |
| 187 | MO15-65 | S | R | R | R | R | R | S |
| 188 | MO15-66 | S | S | R | R | S | R | S |
| 189 | MO15-67 | S | R | R | R | R | R | S |
| 190 | MO15-68 | S | R | R | R | R | R | S |
| 191 | MO15-69 | S | R | R | R | R | R | S |
| 192 | MO15-70 | S | S | R | R | S | R | S |
| 193 | MO15-71 | S | R | R | R | R | R | S |
| 194 | MO15-72 | S | S | R | R | S | R | S |
| 195 | MO15-73 | S | S | S | S | S | S | S |
| 196 | MO15-74 | S | S | R | R | S | R | S |
| 197 | MO15-75 | S | S | S | S | S | S | S |
| 198 | MO15-76 | S | S | S | S | S | S | S |
| 199 | MO15-77 | S | R | R | R | R | R | S |
| 200 | MO15-78 | S | S | S | S | S | S | S |
| 201 | MO15-79 | S | S | R | R | S | R | S |
| 202 | MO15-80 | S | S | R | R | S | R | S |
| 203 | MO15-81 | S | R | R | R | R | R | S |
| 204 | MO15-82 | S | S | R | R | S | R | S |
| 205 | MO15-83 | S | S | S | S | S | S | S |
| 206 | MO15-84 | S | S | S | S | S | S | S |
| 207 | MO15-85 | S | R | R | R | R | R | S |
| 208 | MO15-86 | S | R | R | R | R | R | S |
| 209 | MO15-87 | S | S | S | S | S | S | S |
| 210 | MO15-88 | S | S | S | S | S | S | S |
| 211 | MO15-89 | S | S | S | S | S | S | S |
| 212 | MO15-90 | S | S | S | S | S | S | S |
| 213 | MO15-91 | S | R | R | R | R | R | S |
| 214 | MO15-92 | S | S | S | S | S | S | S |
| 215 | MO15-93 | S | S | S | S | S | S | S |

| | | | | | | | | |
|---|---|---|---|---|---|---|---|---|
| 216 | MO15-94 | S | S | S | S | S | S | S |
| 217 | MO15-95 | S | S | S | S | S | S | S |
| 218 | MO15-96 | S | S | S | S | S | S | S |
| 219 | MO15-97 | S | R | R | R | R | R | S |
| 220 | MO15-98 | S | S | S | S | S | S | S |
| 221 | MO15-99 | S | R | R | R | R | R | S |
| 222 | MO15-100 | S | R | R | R | R | R | S |
| 223 | MO15-101 | S | S | S | S | S | S | S |
| 224 | MO15-102 | S | R | R | R | R | R | S |
| 225 | MO15-103 | S | S | S | S | S | S | S |
| 226 | MO15-104 | S | S | S | S | S | S | S |
| 227 | MO15-105 | S | S | S | S | S | S | S |
| 228 | MO15-106 | S | S | S | S | S | S | S |
| 229 | MO15-107 | S | R | R | R | R | R | S |
| 230 | MO15-108 | S | R | R | R | R | R | S |
| 231 | MO15-109 | S | S | R | R | S | R | S |
| 232 | MO15-110 | S | S | S | S | S | S | S |
| 233 | MO15-111 | S | S | S | S | S | S | S |
| 234 | MO15-112 | S | S | R | R | S | R | S |
| 235 | MO15-113 | S | R | R | R | R | R | S |
| 236 | MO15-115 | S | R | R | R | R | R | S |
| 237 | MO15-116 | S | S | S | S | S | S | S |
| 238 | MO15-117 | S | R | R | R | R | R | S |
| 239 | MO15-118 | S | S | R | R | S | R | S |
| 240 | MO15-119 | S | S | S | S | R | S | S |
| 241 | MO15-120 | S | S | S | S | S | S | S |
| 242 | MO15-121 | S | S | R | R | S | R | S |
| 243 | MO15-122 | S | S | R | R | S | R | S |
| 244 | MO15-123 | S | R | R | R | R | R | S |
| 245 | MO15-124 | S | S | S | S | S | S | S |
| 246 | MO15-125 | S | S | S | S | S | S | S |
| 247 | MO15-126 | S | S | S | S | S | S | S |
| 248 | MO15-127 | S | R | R | R | R | R | S |
| 249 | MO15-128 | S | S | R | R | S | R | S |
| 250 | MO15-129 | S | R | R | R | R | R | S |
| 251 | MO15-130 | S | S | R | R | S | R | S |
| 252 | MO15-131 | S | S | S | S | S | S | S |
| 253 | MO15-132 | S | S | R | R | S | R | S |
| 254 | MO15-133 | S | R | R | R | R | R | S |
| 255 | MO15-134 | S | S | S | S | S | S | R |
| 256 | MO15-135 | S | R | R | R | R | R | S |
| 257 | MO15-136 | S | S | S | S | S | S | S |
| 258 | MO15-137 | S | S | S | S | S | S | S |
| 259 | MO15-138 | S | S | S | S | S | S | S |

| | | | | | | | | |
|---|---|---|---|---|---|---|---|---|
| 260 | MO15-139 | S | S | R | R | S | R | S |
| 261 | MO15-140 | S | S | S | S | S | S | S |
| 262 | MO15-141 | S | S | R | R | S | R | S |
| 263 | MO15-142 | S | S | S | S | S | S | S |
| 264 | MO15-143 | S | R | R | R | R | R | S |
| 265 | MO15-144 | S | S | S | S | S | S | S |
| 266 | MO15-145 | S | S | S | S | S | S | S |
| 267 | MO15-146 | S | R | R | R | R | R | S |
| 268 | MO15-147 | S | S | S | S | S | S | S |
| 269 | MO15-148 | S | S | S | S | S | S | S |
| 270 | MO15-149 | S | S | S | S | S | R | S |
| 271 | MO15-150 | S | R | R | R | R | R | S |
| 272 | MO15-151 | S | R | R | R | R | R | S |
| 273 | MO15-153 | S | R | R | R | R | R | S |
| 274 | MO15-154 | S | R | R | R | R | R | S |
| 275 | MO15-155 | S | R | R | R | R | R | S |
| 276 | MO15-156 | S | R | R | R | R | R | S |
| 277 | MO15-157 | S | R | R | R | R | R | S |
| 278 | MO15-158 | S | R | R | R | R | R | S |
| 279 | MO15-159 | S | R | R | R | R | R | R |
| 280 | MO15-160 | S | S | S | R | S | R | S |
| 281 | MO15-161 | S | S | S | R | R | R | S |
| 282 | MO15-162 | S | S | S | S | S | S | S |
| 283 | MO15-163 | S | R | R | R | R | R | S |
| 284 | MO15-164 | S | S | S | S | S | S | S |
| 285 | MO15-165 | S | R | R | R | R | R | S |
| 286 | MO15-166 | S | R | R | R | R | R | S |
| 287 | MO15-167 | S | R | R | R | R | R | S |
| 288 | MO15-168 | S | S | S | R | S | R | S |
| 289 | MO15-169 | S | R | R | R | R | R | S |
| 290 | MO15-170 | S | S | R | S | S | S | S |
| 291 | MO15-171 | S | S | S | R | S | R | S |
| 292 | MO15-172 | S | S | R | S | S | S | S |
| 293 | MO15-174 | S | R | R | R | R | R | S |
| 294 | MO15-175 | S | R | R | R | R | R | S |
| 295 | MO15-176 | S | S | S | S | S | S | S |
| 296 | MO15-177 | S | S | S | S | S | S | S |
| 297 | MO15-178 | S | S | S | S | S | S | S |
| 298 | MO15-179 | S | R | R | R | R | R | S |
| 299 | MO15-180 | S | R | R | R | R | R | S |
| 300 | MO15-181 | S | R | R | R | R | R | S |
| 301 | MO15-182 | S | S | S | S | S | S | S |
| 302 | MO15-183 | S | R | R | R | R | R | S |
| 303 | MO15-184 | S | S | R | R | S | R | S |

| | | | | | | | | |
|---|---|---|---|---|---|---|---|---|
| 304 | MO15-185 | S | R | R | R | R | R | S |
| 305 | MO15-187 | S | R | R | R | R | R | S |
| 306 | MO15-188 | S | S | R | S | S | S | S |
| 307 | MO15-189 | S | S | R | S | S | S | S |
| 308 | MO15-190 | S | S | R | S | S | S | S |
| 309 | MO15-191 | S | S | S | S | S | S | S |
| 310 | MO15-192 | S | R | R | R | R | R | S |
| 311 | MO15-193 | S | S | R | S | S | S | S |
| 312 | MO15-194 | S | S | R | S | S | S | S |
| 313 | MO15-195 | S | S | S | S | S | S | S |
| 314 | MO15-196 | S | R | R | R | R | R | S |
| 315 | MO15-197 | S | S | R | S | S | S | S |
| 316 | MO15-198 | S | R | R | R | R | R | S |
| 317 | MO15-199 | S | R | R | R | R | R | S |
| 318 | MO15-200 | S | S | R | S | S | S | S |
| 319 | MO15-201 | S | S | S | S | S | S | S |
| 320 | MO15-202 | S | S | R | S | S | S | S |
| 321 | MO15-203 | S | R | R | R | R | R | S |
| 322 | MO15-204 | S | R | R | R | R | R | S |
| 323 | MO15-205 | S | R | R | R | R | R | S |
| 324 | MO15-206 | S | S | R | S | S | S | S |
| 325 | MO15-207 | S | S | R | S | S | S | S |
| 326 | MO15-208 | S | R | R | R | R | R | S |
| 327 | MO15-209 | S | S | R | S | S | S | S |
| 328 | MO15-210 | S | S | S | S | S | S | S |
| 329 | MO15-211 | S | R | R | R | R | R | S |
| 330 | MO15-212 | S | R | R | R | R | R | S |
| 331 | MO15-213 | S | S | R | S | S | S | S |
| 332 | MO15-214 | S | R | R | R | R | R | S |
| 333 | MO15-215 | S | R | R | R | R | R | S |
| 334 | MO15-216 | S | R | R | R | R | R | S |
| 335 | MO15-217 | S | R | R | R | R | R | S |
| 336 | MO15-218 | S | R | R | R | R | R | S |
| 337 | MO15-219 | S | R | R | R | R | R | S |
| 338 | MO15-220 | S | R | R | R | R | R | S |
| 339 | MO15-221 | S | R | R | R | R | R | S |
| 340 | MO15-222 | S | S | R | S | S | S | S |
| 341 | MO15-223 | S | S | R | S | S | S | S |
| 342 | MO15-224 | S | R | R | R | R | R | S |
| 343 | MO15-225 | S | S | S | S | S | S | S |
| 344 | MO15-226 | S | R | R | R | R | R | S |
| 345 | MO15-227 | S | R | R | R | R | R | S |
| 346 | MO15-228 | S | R | R | R | R | R | S |
| 347 | MO15-229 | S | S | R | S | S | S | S |

| | | | | | | | | |
|---|---|---|---|---|---|---|---|---|
| 348 | MO15-230 | S | S | R | S | S | S | S |
| 349 | MO15-231 | S | R | R | R | R | R | S |
| 350 | MO15-232 | S | R | R | R | R | R | S |
| 351 | MO15-233 | S | S | R | S | S | S | S |
| 352 | MO15-234 | S | R | R | R | R | R | S |
| 353 | MO15-235 | S | R | R | R | R | R | S |
| 354 | MO15-236 | S | R | R | R | R | R | S |
| 355 | MO15-237 | S | R | R | R | R | R | S |
| 356 | MO15-238 | S | S | S | S | S | S | S |
| 357 | MO15-240 | S | R | R | R | R | R | S |
| 358 | MO15-241 | S | R | R | R | R | R | R |
| 359 | MO15-242 | S | S | R | S | S | S | S |
| 360 | MO15-243 | S | S | R | S | S | S | S |
| 361 | MO15-244 | S | R | R | R | R | R | S |
| 362 | MO15-245 | S | S | S | S | S | S | S |
| 363 | MO15-246 | S | S | R | S | S | S | S |
| 364 | MO15-247 | S | S | S | S | S | S | S |
| 365 | MO15-248 | S | S | S | S | S | S | S |
| 366 | MO15-249 | S | S | S | S | S | S | S |
| 367 | MO15-250 | S | S | R | S | S | S | S |

R, resistant; S, susceptible

Table S2. Primers used in this study

| Primer | Sequence (5'-3') | Application |
|---|---|---|
| AvrPik-In-F | TGCGTGTTACCACTTTTAAC | *AvrPik* knout out transformants screen |
| AvrPik-In-R | TAAAAGCCGGGCCTTTTTTTCC | |
| AvrPik-out-F | TGGAGTCGGGCTCCTCTTTGCG | |
| Hphcon-R | TACTTCTACACAGCCATCGGTC | |
| RGA4-F3 | GGAAAGCTGATATGTTGTCG | Co-segregation analysis of *Piks* |
| RGA4-R3 | ACTCGGAGTCGGAGAGTCAG | |
| Piks-RGA4-F | TGACGTCGTGAAGAAAGAAG | Detection *of Piks*-1 |
| Piks-RGA4-R | TGCTTAACGTAATTCAACAG | |
| Piks-RGA5-F | ATCATTGAAATGCGGTTTAAAG | Detection of *Piks*-2 |
| Piks-RGA5-R | AGACCGGCAACAGGATTCAATC | |
| BZG17-28F | TGTTGAGGAGGAAAGCTGAT | *Piks*-1 CRISPR mutant identification |
| BZG17-28R | TGCAAGTACGGTTGGAGTG | |
| BZG17-29F | AAAGATAGCAACAAAGTAGACG | *Piks*-2 CRISPR mutant identification |
| BZG17-29R | TTCGATTGATTTGGACATG | |
| AvrPik-F | TAAGGCGGACCTCTCGATTC | Detection of *AvrPik* in field isolate |
| AvrPik-R | CATCCACTTTTCTCGCTGTTC | |
| KP3-F2 | TTTTGGCGCGCCGCCCTTGTTCTCTGTACTTCTCACTGC | Piks-1 expression vector construction |
| KP3-R | TTTTGGCGCGCCGACCACTGACCCAACTTGAAAGACTGG | |
| KP4-F | TTTTGGCGCGCCGCAAGATCAGTACCATCACGAGTAATAGCA | Piks-2 expression vector construction |
| KP4-R | TTTTGGCGCGCCAGGCGGGACCGTAGCAAGTCGTGCTGC | |
| Mutant229-F | CATCACCGACCACCACAACCTGGTCTCTT | Generate of Piks-1 mutant at amino acid position 229 |
| Mutant229-R | CAATCGCCGGTGACCTAAGAGACCAGGTTGT | |
| Mutant261-F | CTCCTTCACATCTTCCTTTACTTGGCT | Generate of Piks-1 mutant at amino acid position 261 |
| Mutant261-R | GTTTCTGGAGGTCAGCCAAGTAAAGGAA | |

Table S3. The various interactions and phenotypes between *Piks*-1, *Piks* mutants and *AvrPik* variants in this study.

| Interaction | *AvrPik* variants | *Pik* allelic gene | | | |
|---|---|---|---|---|---|
| | | *Pikm*-1 | *Piks*-$1^{E229Q}$/ *Pikm*-$1^{V261A}$ | *Piks*-$1^{A261V}$/ *Pikm*-$1^{Q229E}$ | *Piks*-1 |
| Interaction in Y2H | AvrPik-A | +++ | – | – | – |
| | AvrPik-C | – | – | – | – |
| | AvrPik-D | +++ | +++ | +++ | – |
| | AvrPik-E | +++ | – | ++ | – |
| Recognition in rice plants | AvrPik-A | +++ | – | – | – |
| | AvrPik-C | – | – | – | – |
| | AvrPik-D | +++ | +++ | – | – |
| | AvrPik-E | +++ | – | – | – |
| Binding *in vitro* (GF) | AvrPik-A | +++ | – | – | – |
| | AvrPik-C | – | – | – | – |
| | AvrPik-D | +++ | +++ | +++ | – |
| | AvrPik-E | +++ | – | – | – |
| CD response in *Nicotiana benthamiana* | AvrPik-A | +++ | – | – | – |
| | AvrPik-C | – | – | – | – |
| | AvrPik-D | +++ | + | + | – |
| | AvrPik-E | +++ | – | – | – |

Y2H, yeast-2-hybrid; GF, Gel filtration; CD, cell death. Y2H and GF interactions used the isolated HMA domains, and in planta experiments were performed with full-length proteins. Recognition in rice plant *Pikm* is rice cv. IRBLKm-Ts. Recognition in rice plant *Piks* and *Piks* mutants is transgenic rice plants.
